# A novel adhesive complex at the base of intestinal microvilli

**DOI:** 10.1101/2021.01.25.428038

**Authors:** Christian Hartmann, Eva-Maria Thüring, Birgitta E. Michels, Denise Pajonczyk, Sophia Leußink, Lilo Greune, Frauke Brinkmann, Mark Glaesner-Ebnet, Eva Wardelmann, Thomas Zobel, M. Alexander Schmidt, Volker Gerke, Klaus Ebnet

## Abstract

Intestinal epithelial cells form dense arrays of microvilli at the apical membrane to enhance their functional capacity. Microvilli contain a protocadherin-based intermicrovillar adhesion complex localized at their tips which regulates microvillar length and packaging. Here, we identify a second adhesive complex in microvilli of intestinal epithelial cells. This complex is localized at the basal region of microvilli and consists of the adhesion molecule TMIGD1, the phosphoprotein EBP50 and the F-actin – plasma membrane cross-linking protein ezrin. Ternary complex formation requires unmasking of the EBP50 PDZ domains by ezrin binding and is strongly enhanced upon mutating Ser162 located in PDZ domain 2 of EBP50. Dephosphorylation of EBP50 at S162 is mediated by PP1α, a serine/threonine phosphatase localized at the microvillar base and involved in ezrin phosphocycling. Importantly, the binding of EBP50 to TMIGD1 enhances the dynamic turnover of EBP50 at microvilli in a Ser162 phosphorylation-dependent manner. We identify an adhesive complex at the microvillar base and propose a potential mechanism that regulates microvillar dynamics in enterocytes.

## Introduction

Intestinal epithelial cells are highly polarized with an apical domain facing the intestinal lumen and a bounded baso-lateral domain that is in contact with adjacent cells and the extracellular matrix. One prominent feature of these cells is a densely packed array of microvilli at their apical domain, collectively known as brush border (BB) (Crawley *et al*, 2014a; Delacour *et al*, 2016; Sauvanet *et al*, 2015b). Microvilli are actin-based structures consisting of ∼20-30 actin filaments that are bundled by actin cross-linking proteins like villin, espin and fimbrin (Crawley *et al*., 2014a; Sauvanet *et al*., 2015b). The bundling of the actin filaments is thought to be necessary to generate and focus the forces required for membrane deformation during microvilli formation, which is mediated by actin polymerization at the barbed ends of actin filaments (Claessens *et al*, 2006; Mooseker *et al*, 1982).

Microvilli formation is a dynamic process (Gorelik *et al*, 2003; Klingner *et al*, 2014; Meenderink *et al*, 2019; Stidwill *et al*, 1984). When the BB forms, microvilli are highly motile and initially form sparse clusters (Crawley *et al*, 2014b; Meenderink *et al*., 2019). Actin assembly at the barbed ends of the core bundles combined with F-actin treadmilling renders microvilli motile and drives lateral collisions between clusters to form larger clusters which further grow in size and eventually form a BB with microvilli that are maximally packed and uniform in length (Crawley *et al*., 2014b; Meenderink *et al*., 2019). Both microvilli clustering during BB formation as well as maximal packing and length uniformity depend on intermicrovillar adhesion mediated by a protocadherin-based adhesive complex which is specifically enriched at the tips of microvilli (intermicrovillar adhesion complex, IMAC). The IMAC is based on trans-heterophilic interaction of protocadherins CDHR2 and CDHR5 which are linked to the underlying actin cytoskeleton through the PDZ domain scaffolding protein USH1C/Harmonin, its binding partner ANKS4B, and the unconventional myosin MYO7B (Crawley *et al*., 2014b; Crawley *et al*, 2016). Loss of function of the IMAC results in severe defects in brush border morphology, like a loss of microvillar clustering, reduced microvillar density and increased length variability (Crawley *et al*., 2014b; Crawley *et al*., 2016; Pinette *et al*, 2019; Weck *et al*, 2016).

After the establishment of a densely packed BB, microvilli dynamics remain high. In cultured kidney epithelial cells, microvilli undergo phases of growth, steady state, and retraction with an estimated average life cycle of 12.1 ± 5.6 min (Gorelik *et al*., 2003; Loomis *et al*, 2003). Microvilli dynamics is regulated by ezrin, an actin filament - plasma membrane cross-linking protein ezrin (Bretscher, 1983; Zwaenepoel *et al*, 2012). Ezrin switches between a closed and an open conformation, regulated by phosphorylation at T567 (Matsui *et al*, 1999). Phosphorylation T567 of ezrin (Matsui *et al*., 1999) occurs specifically at the distal tips of microvilli (Hanono *et al*, 2006; Pelaseyed & Bretscher, 2018; Viswanatha *et al*, 2012), the site of active microvillar growth (Crawley *et al*., 2014a; Sauvanet *et al*., 2015b). Ezrin T567 phosphocycling is required for apical membrane localization of ezrin as well as microvilli formation (Garbett & Bretscher, 2012; Viswanatha *et al*., 2012). How ezrin T567 phosphorylation is restricted to the microvillar tips is poorly understood. Active ezrin interacts with EBP50/NHERF1 (Reczek *et al*, 1997; Weinman *et al*, 1995), a PDZ domain-containing scaffolding protein which binds PP1α (Kremer *et al*, 2015; Zhang *et al*, 2019), the phosphatase predicted to dephosphorylate ezrin at T567 at the microvillar base (Canals *et al*, 2012; Viswanatha *et al*, 2014; Viswanatha *et al*., 2012).

In this study, we identify a novel adhesive complex at the base of microvilli. This complex consists of the immunoglobulin superfamily (IgSF) member Transmembrane and Immunoglobulin Domain-containing Protein 1 (TMIGD1), EBP50, and ezrin. TMIGD1 directly interacts with EBP50 through a PDZ domain-mediated interaction. This interaction requires the open conformation of EBP50 induced by ezrin binding. PP1α-mediated dephosphorylation of EBP50 at S162 localized in the carboxylate binding loop of PDZ2 promotes the interaction of EBP50 with TMIGD1 and enhances the dynamic turnover of EBP50. Our results identify a new adhesive complex localized at the base of microvilli and provide novel insights into the mechanisms which regulate ezrin phosphocycling and microvilli dynamics.

## Results

### TMIGD1 is a novel component of the brush border in intestinal epithelial cells

TMIGD1 is predominantly expressed by epithelial cells of the kidney (Arafa *et al*, 2015; Hartmann *et al*, 2020) and of the small intestine (Cattaneo *et al*, 2011; Zabana *et al*, 2020). In two colon-derived cell lines, i.e. Caco-2_BBe_ cells (Peterson & Mooseker, 1992) and T84 cells (Murakami & Masui, 1980) we observed in a fraction of cells an enrichment of TMIGD1 at the apical membrane domain with a staining pattern strongly reminiscent of microvillar localization (Fig. 1A, Suppl. Fig.1). Given this rather unusual localization for an adhesion receptor we performed co-stainings with various microvillar markers. TMIGD1 co-localized with villin and ezrin but showed only little co-localization with Eps8, a microvillar tip-localized F-actin capping and bundling protein (Zwaenepoel *et al*., 2012) (Fig. 1B). Immunogold electron microscopy (EM) confirmed that TMIGD1 is predominantly localized at the subapical region of microvilli (Fig. 1C) and mostly absent from the distal tips, which have been estimated to span approximately 25% of microvilli in Caco-2_BBe_ cells (Weck *et al*., 2016). We also analyzed LS174T-W4 cells, a human colon-derived cell line in which a polarized brush border at the single cell level can be induced by doxycycline-regulated expression of the adapter protein STRAD resulting in activation of the polarity kinase LKB1 (Baas *et al*, 2004). Upon LKB1 activation TMIGD1 was efficiently recruited from the cytoplasm to the brush border, where it showed partial co-localization with ezrin and weak co-localization with Eps8 (Fig. 1D-F). In organoids derived from intestinal crypts TMIGD1 co-localized with villin at the apical membrane domain (Fig. 1G). Together, these findings identified the adhesion molecule TMIGD1 as a novel component of the intestinal brush border localized at the subapical region of microvilli.

**Figure 1:**
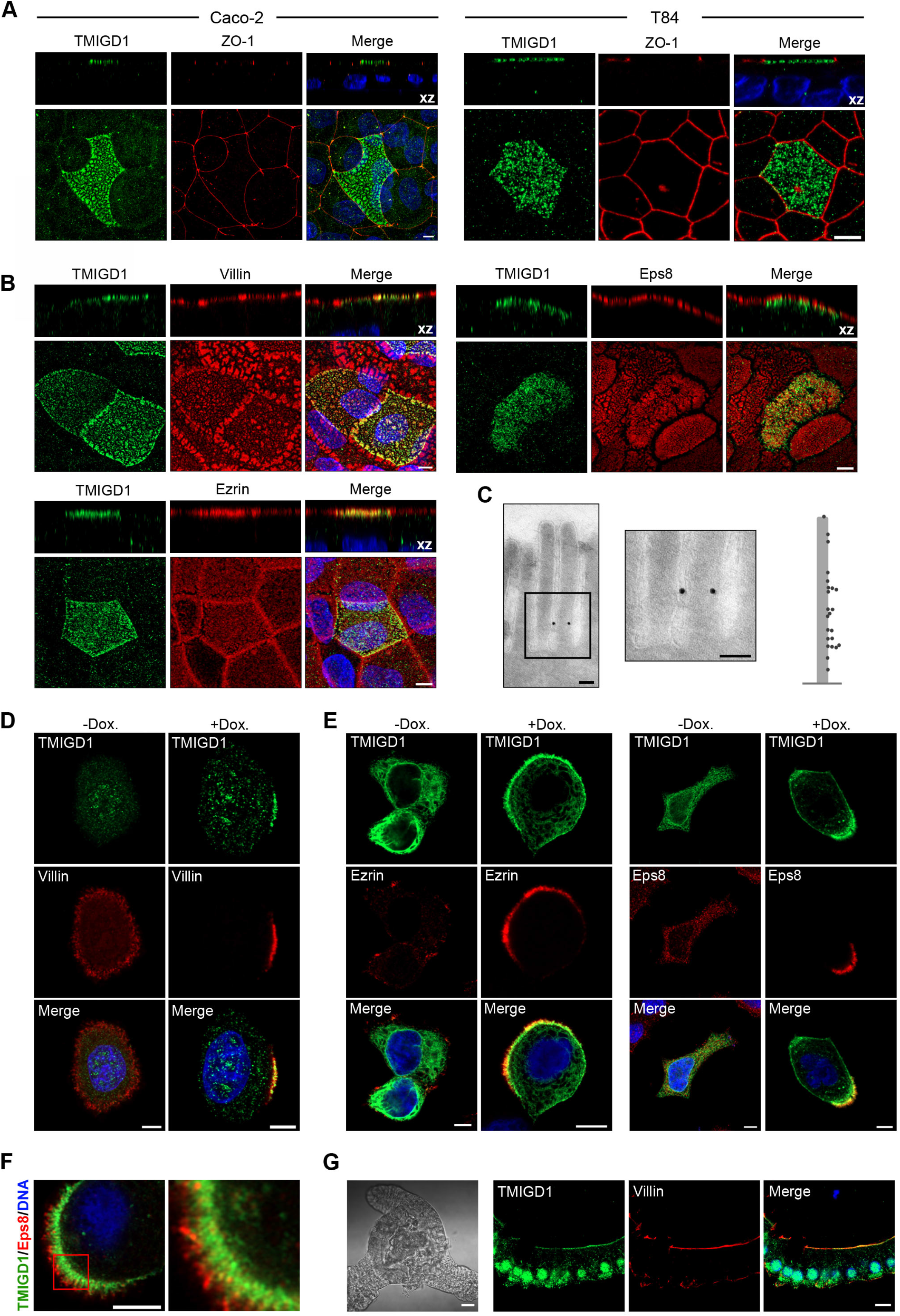
TMIGD1 is a novel component of microvilli. (**A**) IF staining of TMIGD1 in differentiated Caco-2_Bbe1_ cells (left panel) and T84 cells (right panel). Scale bars: 5 µm. (**B**) IF stainings of TMIGD1 with microvilli markers villin, ezrin and Eps8. Scale bars: 5 µm. (**C**) Immunogold TEM of TMIGD1 in differentiated Caco-2_Bbe1_ cells. Middle panel: magnification of the framed area in the left panel. Right panel: Relative position of the gold particles along the microvillar axis. Total number of TMIGD1-positive cells: 23 (6.8% of analyzed cells (n=340). Distribution of gold particles: distal tip: 3; basal region: 20. Scale bar: 100 nm. (**D**) IF stainings of TMIGD1 and villin in LS174T-W4 cells without and with doxycycline (Dox) treatment. Scale bars: 5 µm. (**E**) IF staining of ectopic TMIGD1 and either ezrin or Eps8 in unpolarized (-Dox) and polarized (+Dox) LS174T-W4 cells. Scale bars: 5 µm. (**F**) IF staining of ectopic TMIGD1 and Eps8 in polarized LS174T-W4 cells. Note the localization of TMIGD1 at the subapical region of the brush border and its absence from microvillar distal tips marked by Eps8. Scale bar: 5 µm. (**G**) IF staining of TMIGD1 and villin in murine intestinal organoids. Scale bars: Phase contrast image: 20 µm; IF stainings: 10 µm.

### TMIGD1 directly interacts with two scaffolding proteins localized at the brush border

We next sought to identify cytoplasmic interaction partners for TMIGD1. In a yeast-two hybrid screen we isolated a cDNA fragment that covered AA 140-307 of murine E3KARP/NHERF2. The isolated cDNA clone comprised the entire PDZ2 domain of E3KARP (AA 151-231 of murine E3KARP) (Fig. 2A). *In vitro* binding experiments using the cytoplasmic tail of TMIGD1 fused to GST and an *in vitro* translated E3KARP construct indicated a direct interaction of TMIGD1 and E3KARP, which was abrogated after deleting the PDZ-binding motif (PBM) at the C-terminus of TMIGD1 (Fig. 2B). Mutating the canonical GLGF motif (Doyle *et al*, 1996) in either of the two PDZ domains of E3KARP (PDZ1: G_20_YGF_23_ > GYAA; PDZ2: G_160_YGF_163_ > GYAA) strongly reduced the interaction, suggesting that both PDZ domains can bind TMIGD1 (Fig. 2C). Co-immunoprecipitation (CoIP) experiments from transfected HEK293T cells indicated that TMIGD1 and E3KARP interact in cells and confirmed that this interaction is mediated by the PBM of TMIGD1 and both PDZ domains of E3KARP (Fig. 2D).

**Figure 2:**
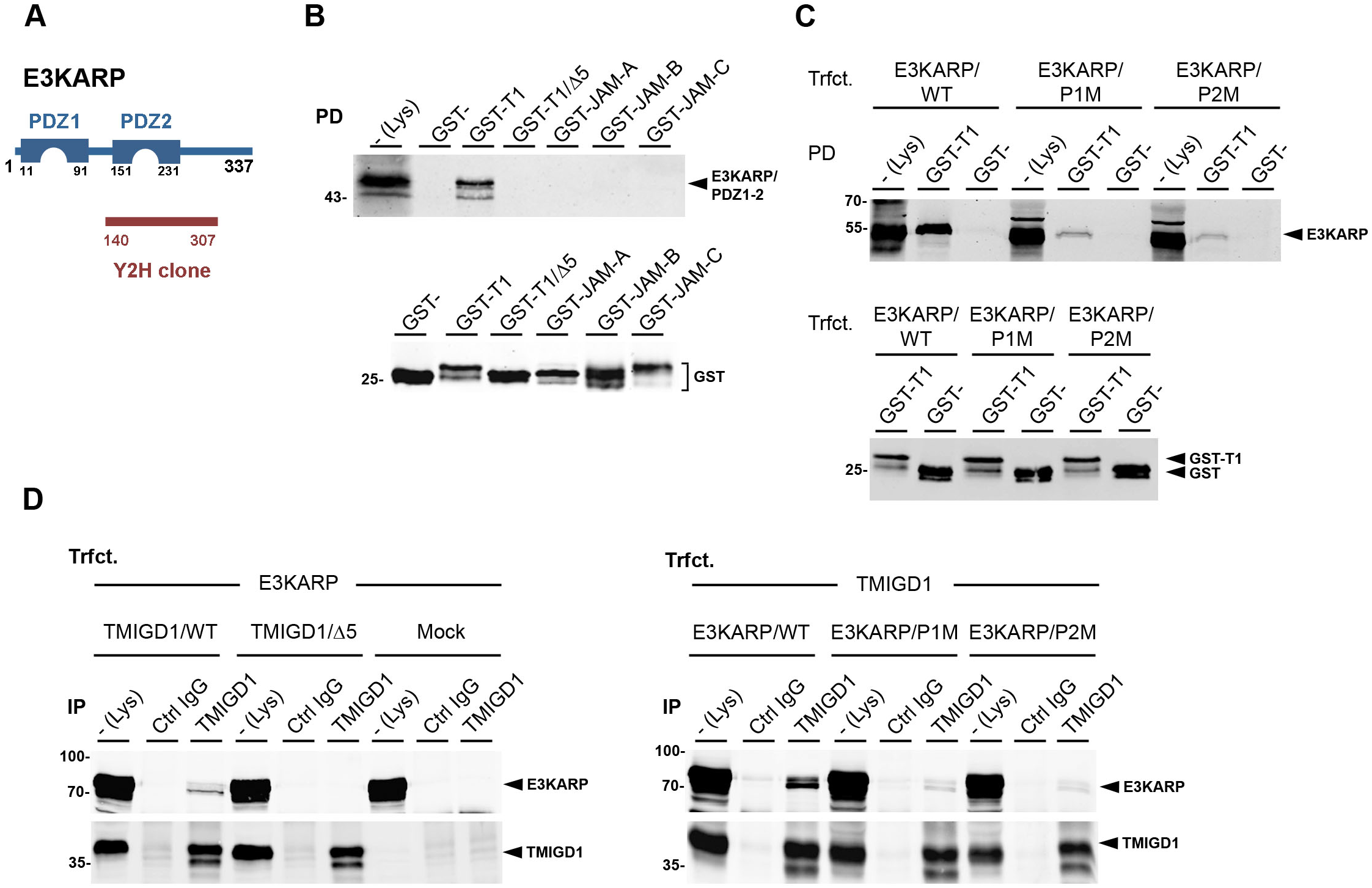
TMIGD1 directly interacts with microvillar scaffolding protein E3KARP. (**A**) Schematic organization of E3KARP. The region isolated as a cDNA clone from a Y2H library interacting with TMIGD1 is depicted in red. (**B**) GST pulldown experiment using the cytoplasmic tails of TMIGD1 (GST-T1, GST-T1/Δ5), JAM-A (GST-JAM-A), JAM-B (GST-JAM-B) or JAM-C (GST-JAM-C) fused to GST and *in vitro* translated E3KARP/PDZ1-2. (**C**) GST-pulldown experiment using GST fusion proteins described in panel B and lysates from HEK293T cells transfected with E3KARP/WT or E3KARP mutants with inactivated PDZ domain 1 (E3KARP/P1M) or PDZ domain 2 (E3KARP/P2M). (**D**) CoIP experiment from HEK293T cells transfected with E3KARP and TMIGD1 deletion constructs (left panel) or transfected with TMIGD1 and E3KARP PDZ domain mutant constructs (right panel).

The closest homologue of E3KARP is EBP50 (or NHERF1) (Reczek *et al*., 1997; Sauvanet *et al*, 2015a). Similar to E3KARP, EBP50 consists of two PDZ domains and a C-terminal ezrin-binding domain (EBD), is localized at microvilli, and has been implicated in microvilli formation (Garbett & Bretscher, 2012; Garbett *et al*, 2010; Garbett *et al*, 2013; LaLonde *et al*, 2010; Morales *et al*, 2004). To test if TMIGD1 also interacts with EBP50, we performed CoIP experiments. EBP50 co-immunoprecipitated with TMIGD1 but not with TMIGD1 lacking the PBM (Fig. 3A). Mutating the PDZ domains in EBP50 (PDZ1: G_23_YGF_26_ > GYAA; PDZ2: G_163_YGF_166_ > GYAA) almost abolished the interaction in both cases (Fig. 3A). To test if this interaction is direct we performed *in vitro* binding experiments with recombinant proteins. Since EBP50 is autoinhibited in the absence of ezrin (Cheng *et al*, 2009; Morales *et al*, 2007; Reczek & Bretscher, 1998), we performed GST pulldown experiments in the presence of recombinant ezrin, either wildtype (dormant) ezrin (Ezrin/WT) or the constitutively active (open) form of ezrin (Ezrin/T567D). In the presence of Ezrin/WT, EBP50 was not detectable in GST-TMIGD1 precipitates (Fig. 3B). However, in the presence of active ezrin (Ezrin/T567D), EBP50 readily co-precipitated with GST-TMIGD1 (Fig. 3B). Mutating either of the two EBP50 PDZ domains strongly reduced the interaction with TMIGD1 (Fig. 3C). Together, these observations indicate that TMIGD1 directly interacts with EBP50 in a PDZ domain-dependent manner, that both PDZ domains of EBP50 can bind TMIGD1, and that this interactions requires the release of EBP50 autoinhibition by active ezrin (Fig. 3D).

**Figure 3:**
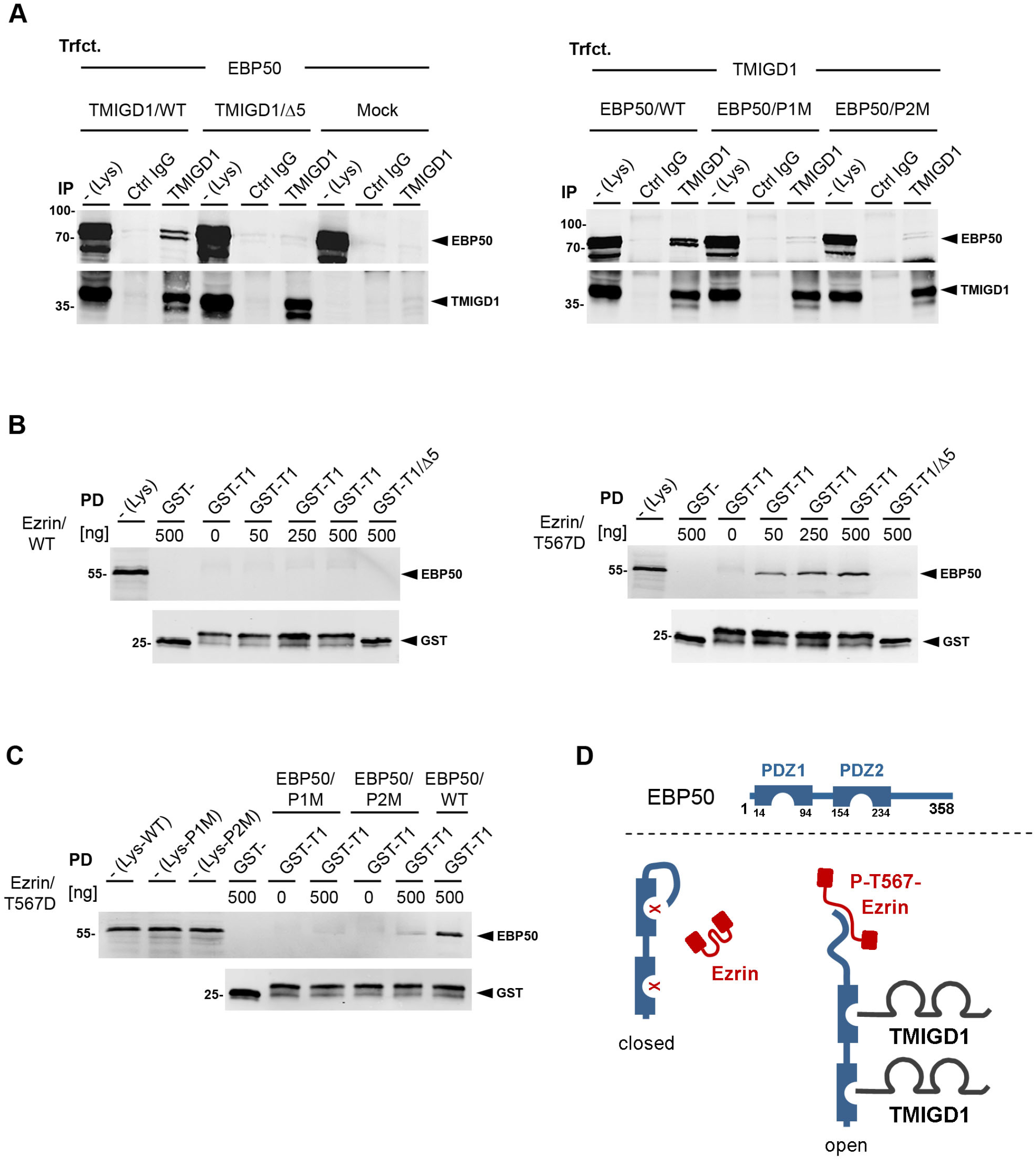
TMIGD1 directly interacts with microvillar scaffolding protein EBP50 in an ezrin-regulated manner. (**A**) CoIP experiment from HEK293T cells transfected with EBP50 and TMIGD1 constructs (TMIGD1/WT, TMIGD1/Δ5, left panel), or transfected with TMIGD1 and EBP50 PDZ domain mutant constructs (PDZ 1 mutant (EBP50/P1M), PDZ 2 mutant (EBP50/P2M) (right panel). (**B**) GST pulldown with GST-TMIGD1 cytoplasmic tail fusion proteins (GST-T1, GST-T1/Δ5) and *in vitro* translated EBP50 in the presence of recombinant wildtype ezrin (left panel) or recombinant ezrin/T567D (right panel). (**C**) GST pulldown with GST-TMIGD1 cytoplasmic tail fusion proteins (GST-T1) and *in vitro* translated EBP50 PDZ domain mutant constructs (EBP50/P1M, EBP50/P2M). (**D**) Schematic depicting the interaction between TMIGD1 and EBP50 and its regulation by active ezrin.

### TMIGD1 is recruited to the brush border by both EBP50 and E3KARP

We next tested if EBP50 and/or E3KARP regulate the localization of TMIGD1 at microvilli. Both EBP50 and E3KARP co-localized with TMIGD1 at polarized brush borders in LS174T-W4 cells (Suppl. Fig. S2A). TMIGD1 localization at the brush border was lost after deletion of the PBM of TMIGD1 (Fig. 4A) indicating a PDZ domain-dependent recruitment to microvilli. Knockdown of either EBP50 or E3KARP alone had only little effect on TMIGD1 recruitment to polarized brush borders (Fig. 4B). Simultaneous knockdown of EBP50 and E3KARP, however, completely abrogated TMIGD1 recruitment to polarized brush borders (Fig. 4B). The combined absence of EBP50 and E3KARP did not change the expression of microvillar key components like ezrin or Eps8 (Supl. Fig. S2B) nor did it prevent the formation of microvilli-containing, F-actin-rich and Eps8-positive polarized caps (Supl. Fig. S2C, D). These findings indicate that both EBP50 and E3KARP can recruit TMIGD1, and that one of these two scaffolding proteins is necessary for microvillar localization of TMIGD1.

**Figure 4:**
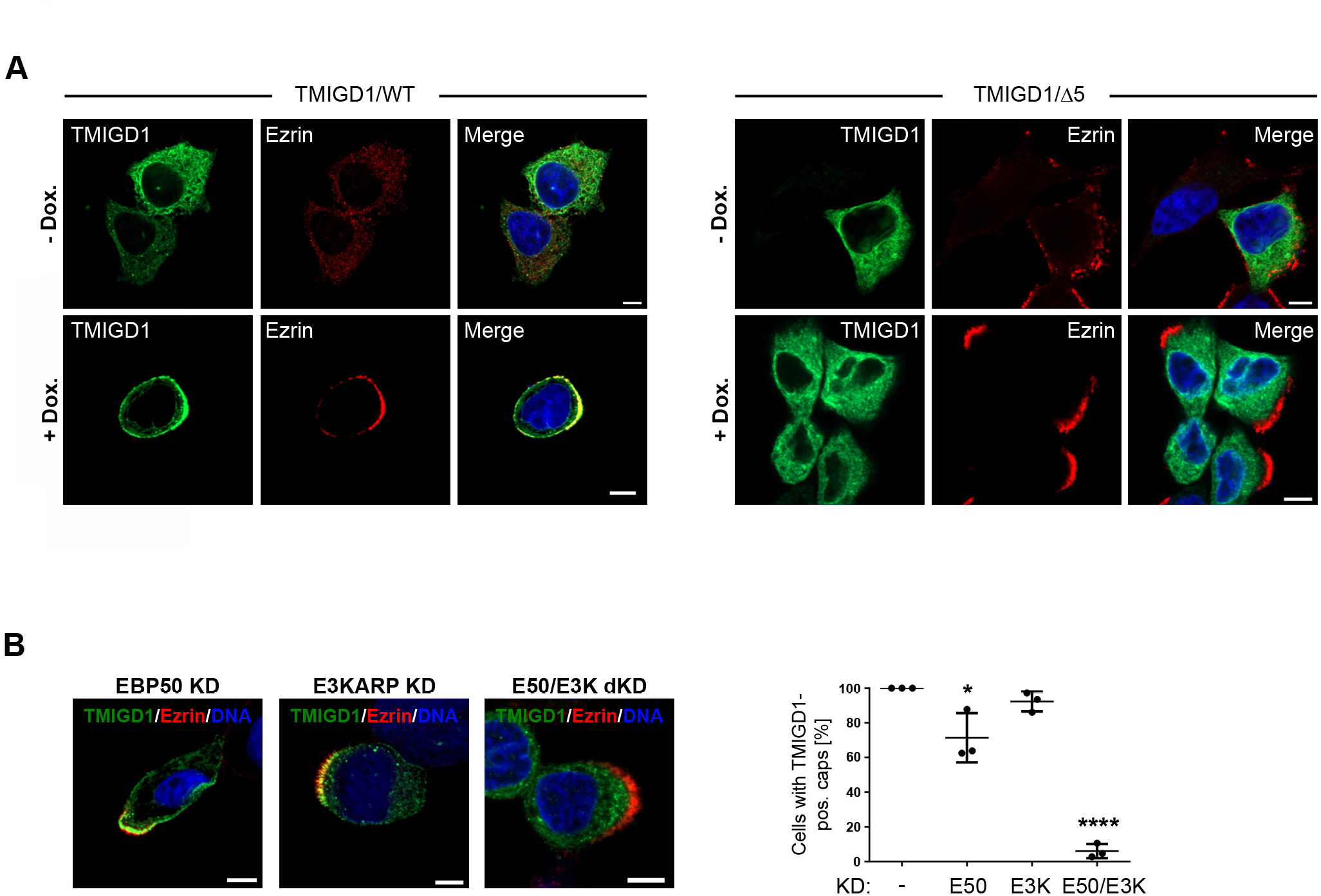
TMIGD1 is recruited to the brush border by EBP50 and E3KARP. (**A**) IF staining of transfected TMIGD1/WT (left panel) and TMIGD1/Δ5 (right panel) in unpolarized (-Dox) and polarized (+Dox) LS174T-W4 cells. Scale bars: 5 µm. (**B**) Left panel: Representative IF images showing the localization of TMIGD1 at polarized brush borders in EBP50 KD, E3KARP KD and EBP50/E3KARP double KD (E50/E3K dKD) LS174T-W4 cells. Scale bars: 5 µm. Right panel: Quantitative analysis of cells with TMIGD1 localization at polarized brush borders. Quantification was performed with unpaired Student’s t-test. Data is presented as mean values ± SD (EBP50/WT: n=142; EBP50 KD: n=102; E3KARP KD: n=92; EBP50/E3KARP dKD: n=99, 3 independent experiments). Ezrin localization was used as marker for polarized brush border formation.

### TMIGD1 regulates the dynamic turnover of EBP50 and E3KARP

The growth of microvilli is a dynamic process with microvillar lifetimes in a range of 7 to 15 min (Garbett & Bretscher, 2012; Gorelik *et al*., 2003). Proteins regulating microvilli growth and dynamics have high turnover rates. Half-maximal fluoresence recovery after photobleaching (FRAP) are in a range of 30 sec for microvilli proteins such as ezrin, E3KARP or brush border Myosin I (MYOIA) (Coscoy *et al*, 2002; Garbett & Bretscher, 2012; Garbett *et al*., 2013; Tyska & Mooseker, 2002). The turnover rate of EBP50 is even faster (half-maximal FRAP ∼ 5 sec) and, interestingly, its turnover is regulated by PDZ domain occupancy (Garbett & Bretscher, 2012; Garbett *et al*., 2013). To test if TMIGD1 influences the dynamics of EBP50 and E3KARP at microvilli we performed FRAP experiments in JEG-3 cells expressing TMIGD1 under a doxycycline-regulated promoter (Suppl. Fig. S3). JEG-3 cells, a choriocarcinoma-derived cell line, has widely been used as a model to study microvilli formation and dynamics (Garbett & Bretscher, 2012; Garbett *et al*., 2010; Garbett *et al*., 2013; Hanono *et al*., 2006; Sauvanet *et al*., 2015a; Viswanatha *et al*., 2012). The recovery rates for ezrin did not significantly differ in the absence and presence of TMIGD1 expression (Fig. 5A). The recovery rates of both EBP50 and E3KARP were slightly but significantly reduced in the presence of TMIGD1 expression (EBP50: P=0.031, E3KARP: P=0.026) (Fig. 5B, C). These observations suggest that TMIGD1 reduces the mobility of EBP50 and E3KARP in microvilli, most likely by immobilizing the two proteins at the microvillar membrane.

**Figure 5:**
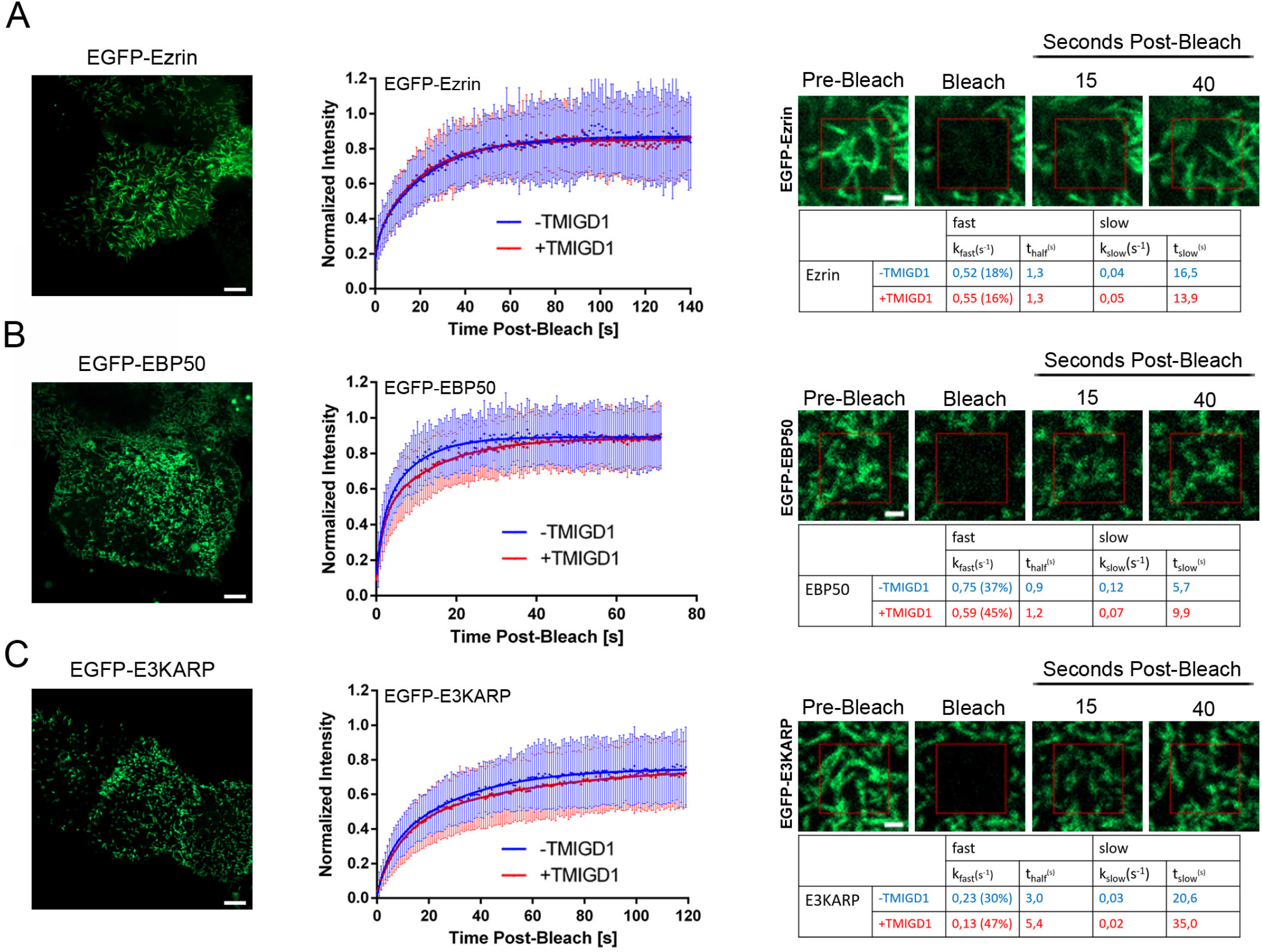
TMIGD1 influences the dynamics of EBP50 and E3KARP. FRAP experiments for ezrin (A), EBP50 (B) and E3KARP (C). Left panels: Representative images of EGFP-Ezrin, EGFP-EBP50 and EGFP-E3KARP in apical microvilli of JEG-3 cells. Scale bars: 5 µm. Middle panels: Photobleaching recovery curves in cells without TMIGD1 expression (-TMIGD1) and cells with doxycycline-induced TMIGD1 expression (+TMIGD1) of EGFP-Ezrin (n=50 and n=62, -TMIGD1 and +TMIGD1, respectively), EGFP-EBP50 (n=81 and n=89, -TMIGD1 and +TMIGD1, respectively), and EGFP-E3KARP (n=143 and n=138, -TMIGD1 and +TMIGD1, respectively). Error bars show mean values ± SD. Statistical analysis was performed using Two-way Repeated Measurements ANOVA. Statistical significance: EGFP-Ezrin, not significant; EGFP-EBP50, P=0.031; EGFP-E3KARP, P=0.026. Right panels: Representative fluorescence images at different time points of FRAP experiments for EGFP-Ezrin, EGFP-EBP50 and EGFP-E3KARP. Tables show recovery rates (k) and half maximal recovery times (t_half_) of double exponential (fast and slow) recovery curves. Scale bars: 1 µm.

### EBP50 phosphorylation at Ser162 regulates TMIGD1 binding and protein turnover

EBP50 is a phosphoprotein with numerous phosphorylation sites (Reczek *et al*., 1997; Vaquero *et al*, 2017). Several phosphoserine residues are implicated in microvilli assembly, including S162, S280 and S302 which are phosphorylated by PKC (Garbett *et al*., 2010; Li *et al*, 2007; Raghuram *et al*, 2003), and S339 and S240 which are phosphorylated by Cdk1 (Garbett *et al*., 2010; He *et al*, 2001). We generated phosphodeficient Ser-to-Ala mutants of these residues in three groups (S162A, S280-302A, S339-340A) and analyzed the interaction of these mutants with TMIGD1 in CoIP experiments. The S280-302A and S339-340A mutations did not alter the interaction with TMIGD1 (Fig. 6A). Surprisingly, the S162A mutation significantly enhanced the interaction with TMIGD1 (Fig. 6A) suggesting that the TMIGD1 - EBP50 interaction is subject to phosphoregulation by PKC involving S162.

**Figure 6:**
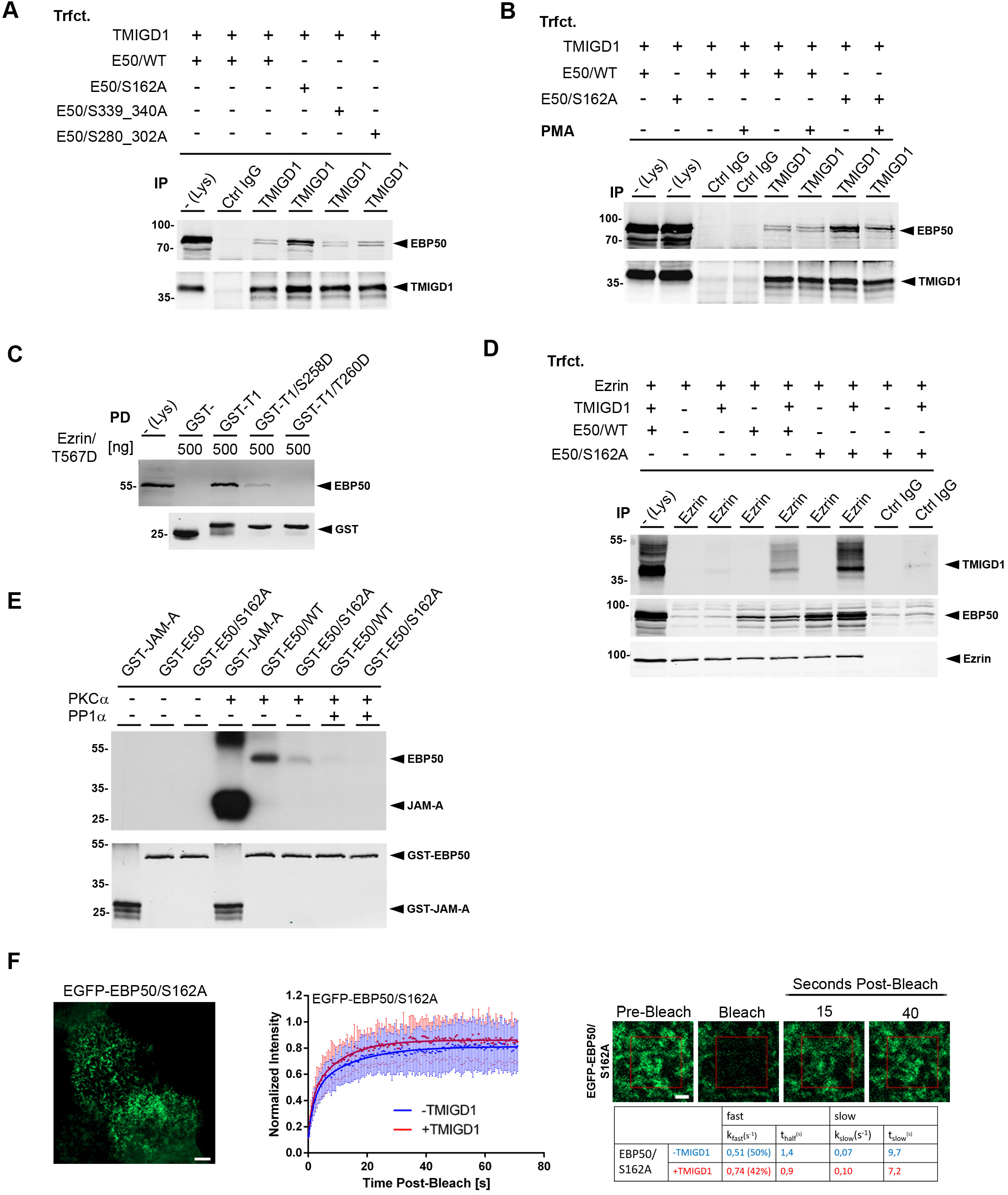
TMIGD1 binding to EBP50 is regulated by phosphorylation of S162 in the carboxylate binding loop of PDZ2. (**A**) CoIP experiment from HEK293T cells transfected with TMIGD1 and various EBP50 phosphodeficient mutant constructs (E50/S162A, E50/S339_340A, E50/S280_302A). Note the strong increase in TMIGD1 binding in the EBP50/S162A mutant. (**B**) CoIP experiment from HEK293T cells transfected with TMIGD1 and EBP50 constructs (E50/WT, E50/S162A) in the presence of PMA. (**C**) GST pulldown with the cytoplasmic tail of TMIGD1, either WT (GST-T1) or mutants mimicking phosphorylation of Ser/Thr residues present in the PDZ domain-binding motif of TMIGD1 (GST-T1/S258D, GST-T1/T260D), fused to GST and *in vitro* translated EBP50 in the presence of recombinant ezrin/T567D. (**D**) CoIP experiment from HEK293T cells transfected with ezrin (the Δ3 mutant of ezrin was used because ezrin/WT interacts only poorly with EBP50 in cells (Viswanatha *et al*., 2013), TMIGD1 and either EBP50/WT (E50/WT) or EBP50/S162A (E50/S162A). Note that TMIGD1 co-immunoprecipitation with ezrin requires EBP50 and that this interaction is enhanced when S162 is mutated. (**E**) *In vitro* phosphorylation/dephosphorylation of EBP50 by PKCα and PP1α. GST-EBP50 fusion proteins (GST-E50/WT (AA 95-268), GST-E50/S162A (AA 95-268, S162A) were incubated with recombinant PKCα either alone or followed by incubation with PP1α in the presence of radiolabeled Y-^32^P-ATP. GST-JAM-A served as positive control for PKCα-mediated phosphorylation. Phosphorylation was analyzed by autoradiography (top panel), equal loading of GST fusion proteins was analyzed by Western blotting (bottom panel). (**F**) FRAP experiments for EBP50/S162A. Left panel: Representative image of EGFP-EBP50/S162A at apical microvilli of JEG-3 cells. Scale bar: 5 µm. Middle panel: Photobleaching recovery curves of EGFP-EBP50/S162A in cells without TMIGD1 expression (-TMIGD1, n=84) and cells with doxycycline-induced TMIGD1 expression (+TMIGD1, n=71). Error bars show mean values ± SD. Statistical analysis was performed using Two-way Repeated Measurements ANOVA. Statistical significance: P=0.0007. Right panels: Representative fluorescence images at different time points of FRAP experiments for EGFP-EBP50/S162A. The table shows recovery rates (k) and half maximal recovery times (t_half_) of double exponential (fast and slow) recovery curves. Scale bar: 1 µm.

To further explore a PKC-dependent interaction between TMIGD1 and EBP50, we stimulated cells with PMA. EBP50/WT co-immunoprecipitated with TMIGD1 at similar efficiencies without or with PMA stimulation (Fig. 6B). EBP50/S162A, however, co-immunoprecipitated with TMIGD1 at much lower efficiencies after PMA stimulation (Fig. 6B). These findings further suggest that the interaction between TMIGD1 and EBP50 is dynamic and subject to PKC-mediated phosphoregulation.

Intriguingly, S162 of EBP50 is located within PDZ domain 2 and is situated in immediate vicinity of the GLGF motif (G_163_YGF_166_) suggesting that its phosphorylation impairs ligand binding. We therefore tested the possibility that TMIGD1 phosphorylation within the PBM could block the interaction with EBP50. We generated phosphomimetic mutants of both potential PKC phosphorylation sites present in the PBM of TMIGD1 (S258D, T260D) and performed *in vitro* binding experiments. Mimicking phosphorylation of either S258 or T260 strongly impaired the interaction with EBP50 (Fig. 6C), suggesting that the interaction of TMIGD1 and EBP50 may be regulated by phosphorylation of amino acids located at the interface between the two interacting molecules.

The function of EBP50 in microvilli formation has been found to depend on its interaction with both ezrin and a hitherto unidentified PDZ1 ligand (Garbett *et al*., 2010), which suggested the existence of an integral membrane protein at microvilli that tethers the ezrin - EBP50 complex to the membrane. We therefore tested if ezrin is associated with TMIGD1 through EBP50. Since EBP50 associates poorly with full length ezrin, most likely because ezrin undergoes an intramolecular head-to-tail interaction involving the N-terminal FERM domain and the C-terminal E_584_A_585_L_586_ motif (Gary & Bretscher, 1995; Viswanatha *et al*, 2013), we used a C-terminal truncation mutant of ezrin (ezrin_1-583_) for these experiments. TMIGD1 co-immunoprecipitated with ezrin when EBP50 was present but not when EBP50 was absent (Fig. 6D). The interaction of TMIGD1 and ezrin was enhanced when S162 of EBP50 was mutated to Ala (Fig. 6D). These findings indicated that TMIGD1 exists in a ternary complex with ezrin and EBP50 and suggest that TMIGD1 tethers ezrin to the plasma membrane through EBP50. They also indicate that dephosphorylation of EBP50 at S162 enhances the tethering of the ezrin – EBP50 complex to the membrane by TMIGD1.

EBP50 and ezrin are localized along the entire microvilli but active ezrin characterized by T567 phosphorylation is restricted to the distal tips (Garbett *et al*., 2013; Hanono *et al*., 2006). Inactivation of ezrin at the basal region has been proposed to involve dephosphorylation of T567 by protein phosphatase PP1α (Canals *et al*., 2012; Viswanatha *et al*., 2014; Viswanatha *et al*., 2012). Intriguingly, PP1α can directly interact with EBP50 through a conserved VxW/F motif in EBP50 (V_257_P_258_F_259_) (Kremer *et al*., 2015; Zhang *et al*., 2019) that does not overlap with the ezrin-binding motif (M_346_-L_358_) (Terawaki *et al*, 2006), suggesting that EBP50 could act as a scaffold for both ezrin and PP1α allowing simultaneous binding and their functional interaction. To address the possibility that PP1α dephosphorylates Ser162 of EBP50, which would promote the interaction with TMIGD1, we performed *in vitro* dephosphorylation experiments with an EBP50 and recombinant PP1α. We used a GST-EBP50 fusion construct that consists of AA95 - 268, thus containing the S162 residue as the only known PKCα phosphorylation site and in addition the PP1α binding motif (V_257_P_258_F_259_). In line with published data (Li *et al*., 2007; Raghuram *et al*., 2003), GST-EBP50/95-268 was strongly phosphorylated by PKCα (Fig. 6E). The S162A mutant was only barley phosphorylated indicating that Ser162 is the only relevant PKCa phosphorylation site in this constuct. Importantly, PP1α completely dephosphorylated S162-phosphorylated EBP50/95-268 (Fig. 6E). These observations identify PP1α as a phosphatase that dephosphorylates EBP50 at S162. They suggest that PP1α localized at the microvillar base might not only serve to dephosphorylate ezrin at T567 to restrict the ezrin phosphocycle to microvillar tips but also to dephosphorylate EBP50 at S162 in PDZ 2 to promote the binding of EBP50 to TMIGD1 and the tethering of the ezrin - EBP50 complex to the membrane.

Previous observations indicate that EBP50 PDZ domain occupation increases the dynamic behaviour of EBP50 at microvilli (Garbett & Bretscher, 2012; Garbett *et al*., 2013). To test if the P-S162 regulation of the TMIGD1 - EBP50 interaction affects the dynamics of EBP50 turnover at microvilli, we performed FRAP experiments with EBP50/S162A in the absence and presence of TMIGD1. Surprisingly, as opposed to its influence on EBP50/WT, TMIGD1 accelerated the recovery of EBP50/S162 after photobleaching (P=0.0007) (Fig. 6F) suggesting that the binding of S162-dephosphorylated EBP50 to TMIGD1 increases its dynamics at microvilli. These findings suggest that TMIGD1 represents an integral protein at the microvillar base with the capacity to increase the turnover of EBP50 by binding to EBP50 PDZ domains, which has previously been postulated to exist (Garbett & Bretscher, 2012). Altogether, these findings suggest a model in which dephosphorylation of EBP50 at S162 by EBP50-bound PP1α promotes the interaction of the ezrin - EBP50 complex with TMIGD1. At the same time, most likely as a result of PP1α-mediated dephosphorylation of ezrin at T567 which results in ezrin inactivation, the ezrin - EBP50 complex is destabilized resulting in the closed EBP50 conformation, reduced EBP50 binding to TMIGD1 and thus increased turnover of EBP50.

### TMIGD1 is a bona fide adhesion receptor

Microvilli contain a protocadherin-based adhesive complex at their tips, the intermicrovillar adhesion complex (IMAC) (Crawley *et al*., 2014b). Since TMIGD1 has been described as adhesion molecule (Arafa *et al*., 2015), we tested its adhesive properties in more detail. Ectopically expressed TMIGD1 increased aggregate formation of HEK293T cells (Fig. 7A), indicating that TMIGD1 can promote homotypic cell-cell adhesion, as oberved before (Arafa *et al*., 2015). To test if TMIGD1 is a homophilic adhesion receptor, we performed CoIP experiments with differentially tagged TMIGD1 constructs. Untagged TMIGD1 was efficiently immunoprecipitated with Flag-EGFP-TMIGD1 (Fig. 7B). Untagged TMIGD1 was also immunprecipitated with a Flag-EGFP-TMIGD1 construct in which the cytoplasmic domain of TMIGD1 was replaced by the cytoplasmic domain of JAM-A (Fig. 7B). Vice versa, an untagged TMIGD1 lacking the complete cytoplasmic domain was efficiently immunoprecipitated with Flag-EGFP-TMIGD1 (Fig. 7B). These findings indicate that TMIGD1 undergoes homophilic interaction through its extracellular domain, either in cis or in trans. To test if this interaction can occur in trans, we performed aggregation assays with fluorescently labelled beads coated with the extracellular domain (ECD) of TMIGD1. TMIGD1-coated beads formed aggregates (Fig. 7C) indicating a trans-homophilic adhesive activity of TMIGD1 that is sufficient to support aggregation, a property that is typical for strong adhesion molecules like cadherins and various IgSF members (Honig & Shapiro, 2020), and that has recently been described for heterophilic protocadherin interactions at the microvillar tips (Crawley *et al*., 2014b). A model incorporating our findings on TMIGD1 into the existing model on the regulation of microvilli dynamics is depicted in Fig. 8.

**Figure 7:**
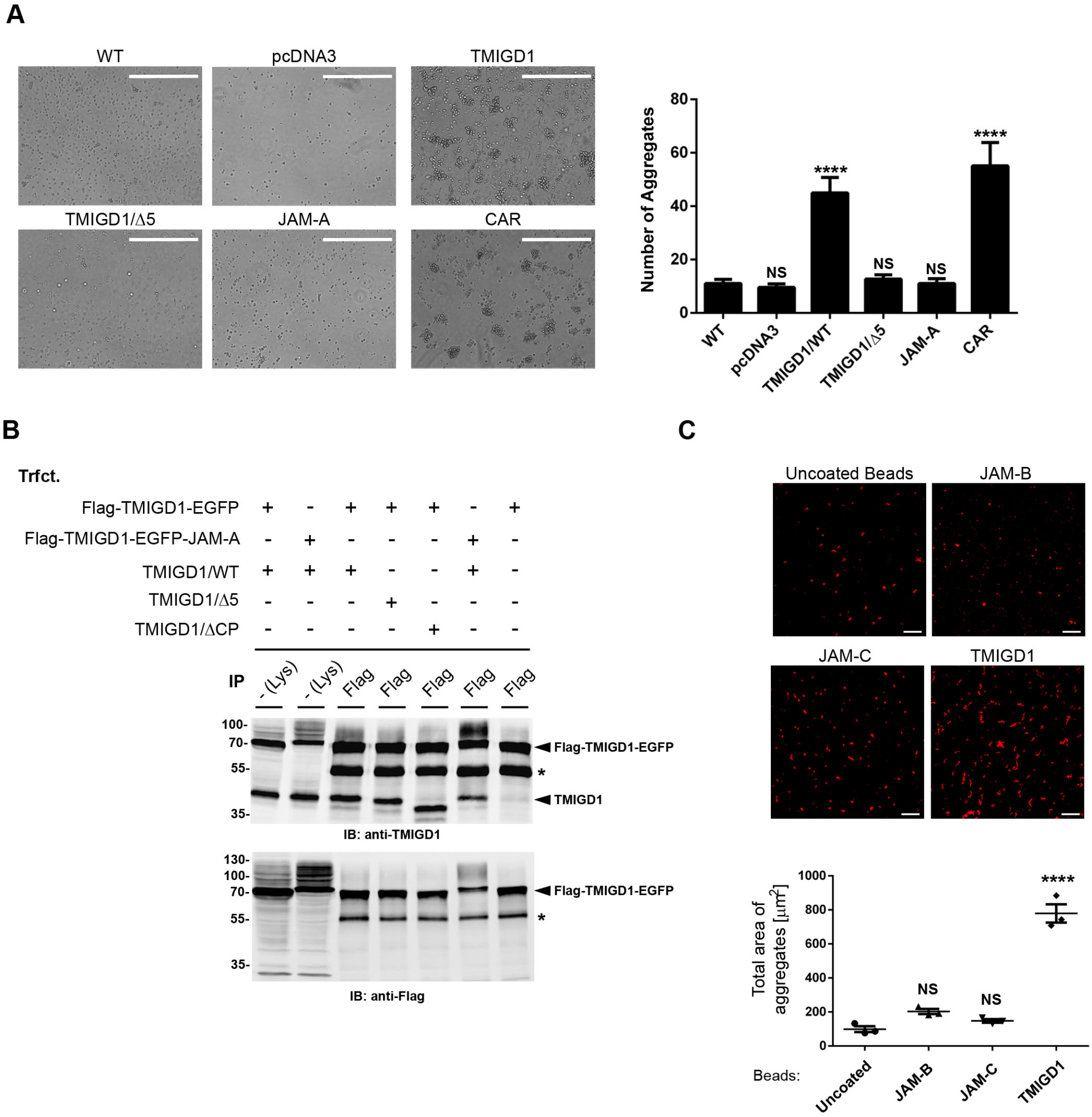
TMIGD1 mediates trans-homophilic interaction. (A) Cell aggregation assays of HEK293T cells either untransfected (WT) or transfected with empty vector (pcDNA3), TMIGD1/WT, TMIGD1/Δ5, JAM-A (negative control) or CAR (positive control). Statistical analysis was performed using unpaired student’s t-test, data is presented as means ± SEM (five independent experiments). (**B**) CoIP experiment from HEK293T cells transfected with untagged TMIGD1 (TMIGD1/WT, TMIGD1/Δ5, TMIGD1/ΔCP, lacks cytoplasmic part) together with tagged TMIGD1 (Flag-TMIGD1-EGFP, Flag-TMIGD1-EGFP-JAM-A (cytoplasmic part of TMIGD1 replaced by cytoplasmic part of JAM-A) as indicated. Asterisks indicate Ig heavy chains. (**C**) Bead aggregation assay with fluorescently labeled beads either uncoated or coated with Fc fusion proteins of JAM-B or JAM-C (negative controls) or TMIGD1. Left panel: Representative IF images. Scale bars: 20 µm. Right panel: Quantification of bead aggregation. Quantification was performed using Ordinary one-way ANOVA with Dunnett’s multiple comparison. Data is presented as means ± SEM (3 independent experiments). ****P<0.0001.

**Figure 8:**
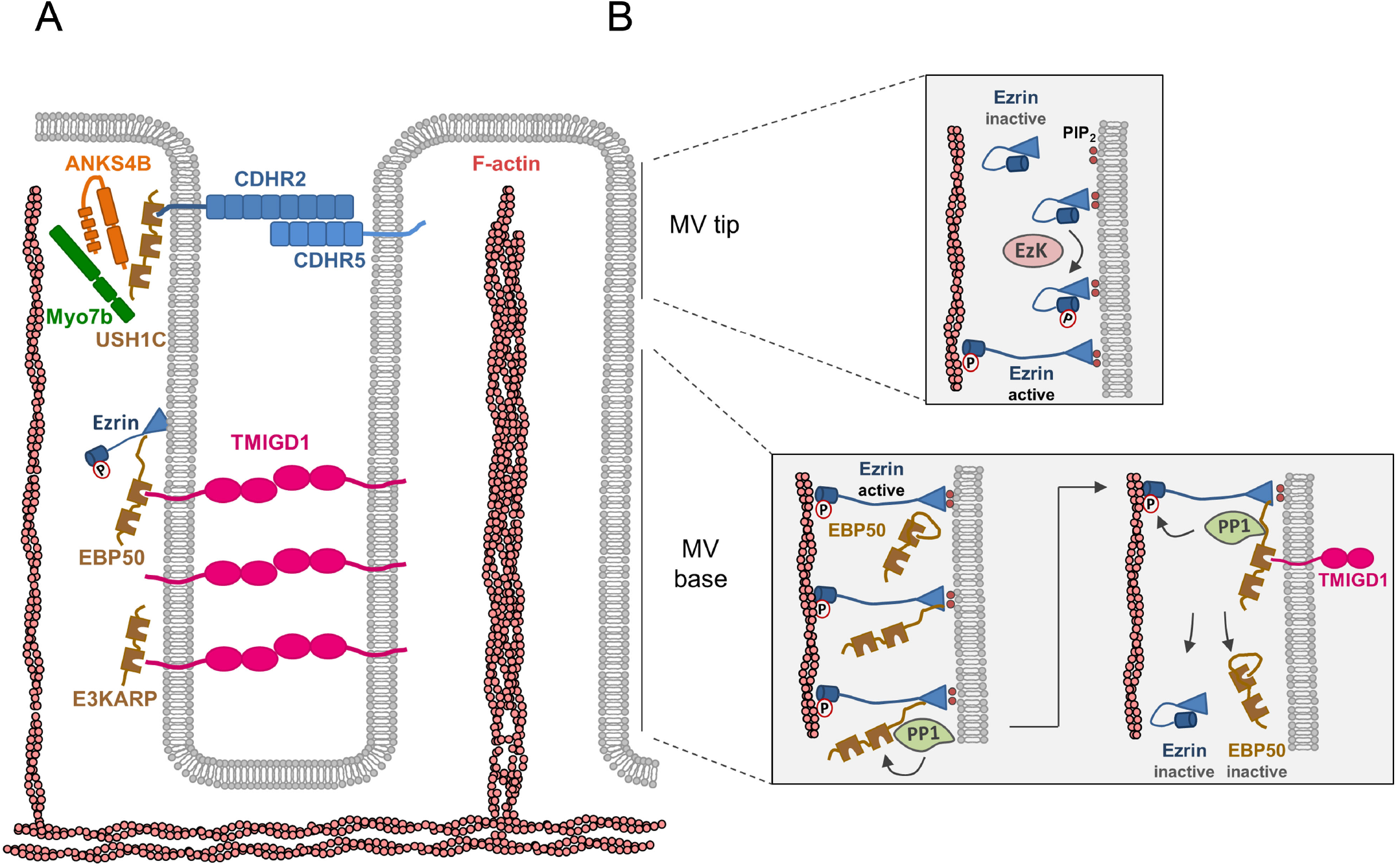
Model of TMIGD1 localization and function at microvilli. (**A**) Intermicrovillar adhesion complexes in epithelial cells. Microvilli (MV) contain a protocadherin-based intermicrovillar adhesion complex (IMAC) at their tips. Protocadherins CDHR2 and CDHR5 interact in a trans-heterophilic manner. MV tip localization of the IMAC is regulated by Myo7B associated with CDHR2 or CDHR5 through USHC1. The IMAC regulates microvilli packing. The present study suggest a second adhesion complex localized at the subapical region of microvilli. This complex is based on trans-homophilic interactions of the IgSF member TMIGD1 which interacts with the two scaffolding proteins EBP50 and E3KARP. (**B**) Hypothetical model on the role of the TMIGD1-based adhesion complex. The following model incorporates the observations presented in this manuscript into the existing model proposed by Viswanatha and colleagues (Viswanatha *et al*., 2014). **MV tip**: Ezrin localized at the MV tips is phosphorylated at T567 by an ezrin kinase (EzK), either lymphocyte oriented kinase (LOK) or the related sterile 20-like kinase (SLK) and/or Mammalian STE20-like protein kinase 4 (MST4). T567-phosphorylated ezrin adopts the open conformation allowing simultaneous binding to the plasma membrane (through the N-terminal FERM domain) and F-actin (through the C-terminal C-ERMAD domain). **MV base**: Through its FERM domain, active ezrin binds to the ezrin-binding region of EBP50 triggering the open conformation of EBP50 thereby enabling accessibility both of its two PDZ domains and of its C-terminal region which harbors the PP1α binding site. Ezrin-bound EBP50 is now able to bind PP1α which dephosphorylates S162 in PDZ domain 2 of EBP50 thereby promoting its interaction with TMIGD1. EBP50-bound PP1α dephosphorylates ezrin at T567 resulting in the closed conformation and ezrin inactivation. As a consequence, ezrin dissociates from EBP50 which adopts its closed conformation and dissociates from TMIGD1. Inactive ezrin and inactive EBP50 can undergo another cycle of activation and deactivation. Adapted from (Viswanatha *et al*., 2014).

## Discussion

In this study we identify a novel adhesion complex in microvilli of enterocytes. The adhesion receptor present in this complex is the IgSF member TMIGD1 which directly interacts with two microvilli-localized PDZ domain-containing scaffolding proteins, EBP50/NHERF1 and E3KARP/NHERF2. The interaction with EBP50 requires ezrin resulting in a ternary complex in which ezrin is linked to membrane-localized TMIGD1 through EBP50. Both ezrin and EBP50 play critical roles in microvilli formation and dynamics (Garbett & Bretscher, 2012; Garbett *et al*., 2010; LaLonde *et al*., 2010; Viswanatha *et al*., 2012). Recent studies described TMIGD1 as component of the apical membrane of enterocytes whose absence results in a defective BB membrane (De La Cena *et al*, 2020; Zabana *et al*., 2020). Our findings thus identify TMIGD1 as part of an adhesive complex at the micorvillar base and describe a mechanism though which TMIGD1 regulates microvilli formation.

TMIGD1 has been described to be predominantly expressed by epithelial cells of the kidney and of the intestine (Arafa *et al*., 2015; Cattaneo *et al*., 2011; Meyer *et al*, 2018). Its expression is downregulated during malignant transformation and during inflammation both in the kidney (Arafa *et al*., 2015; Meyer *et al*., 2018) and in the intestine (Cattaneo *et al*., 2011; Lee *et al*, 2015; Mojica & Hawthorn, 2010; Roberts *et al*, 2015; Zabana *et al*., 2020) suggesting a tumor-suppressive function. The reduction and/or loss of TMIGD1 expression during cellular transformation might explain the low expression levels of TMIGD1 in many cell lines derived from kidney or intestine (Cattaneo *et al*., 2011; Meyer *et al*., 2018) and might also be responsible for our observation of TMIGD1 expression in only a fraction of colon carcinoma-derived Caco-2 and T84 cells (Fig. 1A, Suppl. Fig. S1).

We find that TMIGD1 directly interacts with two scaffolding proteins at microvilli, EBP50 and E3KARP, and that either of the two is required for microvillar localization of TMIGD1. In both cases, the interaction is direct and mediated by the PDZ domain-binding motif of TMIGD1 (-SETAL). We observed for both EBP50 and E3KARP that inactivating either of the two PDZ domains strongly impairs TMIGD1 binding (Fig. 2, 3). We interpret this in a way that both PDZ domains can bind TMIGD1 and that ligand binding to either of the two PDZ domains may influence the accessibility of the other PDZ domain for TMIGD1. In line with this assumption, autoinhibition of EBP50 mediated by its C-terminal PBM (-LFSNL) interacting with PDZ2 masks both PDZ domains (Morales *et al*., 2007), and vice versa, the release of autoinhibition by ezrin binding to the EB region renders both PDZ domains accessible for ligands (Li *et al*, 2009). Also, studies with EBP50 phosphomimicking mutants showed that ligand binding to PDZ2 blocks PDZ1 accessibility (Garbett *et al*., 2010). This coordinated regulation of PDZ domain accessibility is most likely regulated by an allosteric mechanism in which ligand binding induces a long range conformational change of EBP50 (Bhattacharya *et al*, 2010; Bhattacharya *et al*, 2019; Li *et al*., 2009). An interaction of both PDZ domains with the same ligand has also been found for the interaction of EBP50 with the cystic fibrosis transmembrane conductance regulator (CFTR) as well as for the interaction of E3KARP with PTEN (Short *et al*, 1998; Takahashi *et al*, 2006).

The interaction of TMIGD1 with EBP50 is enhanced when S162 of EBP50 is mutated to Ala, strongly suggesting that Ser162 phosphorylation inhibits the interaction. Ser162 is located in PDZ2 of EBP50 and immediately precedes the canonical GLGF motif present in the carboxylate-binding loop of PDZ domains (G_163_YGF_166_ in EBP50 PDZ2) (Doyle *et al*., 1996). Similarly positioned phosphoserine residues with negative impact on ligand binding have been identified in PDZ1 of DLG1/SAP-97 (S_232_GLGF_236_) and in PDZ1 of DLG4/PSD-95 (S_73_GLGF_77_), (Gardoni *et al*, 2006; Mauceri *et al*, 2007; Pedersen *et al*, 2017). In addition, the interaction of EBP50 with the cystic fibrosis transmembrane conductance regulator (CFTR) has been described to be negatively regulated by S162 phosphorylation of EBP50 (Raghuram *et al*., 2003). Site-specific phosphorylation of both PDZ domains and the PBMs in their ligands has been described to fine-tune and to regulate specificity of PDZ domain – ligand interactions (Liu & Fuentes, 2019). As a further evidence for a phosphoregulation of the EBP50 – TMIGD1 interaction, we found that mimicking phosphorylation of two residues present in the PBM of TMIGD1 prevents the interaction with EBP50. Our observations thus suggest that the TMIGD1 - EBP50 interaction is regulated by phosphorylation of residues localized at key positions in the PDZ domain - PDZ ligand interface thus allowing a rapid and dynamic regulation of the interaction.

We find that PKCα-phosphorylated S162 of EBP50 is dephosphorylated by PP1α. EBP50 contains a PP1 binding motif in the linker region between PDZ2 and the EBD and acts as scaffold for PP1α (Kremer *et al*., 2015; Zhang *et al*., 2019). EBP50-bound PP1α dephosphorylates Ser290 of EBP50, a major phosphorylation site for GRK6A (Hall *et al*, 1999; Zhang *et al*., 2019) which, however, is not involved in microvilli formation (Garbett *et al*., 2010). Intriguingly, PP1α has also been postulated to be localized at the basal region of microvilli and to dephosphorylate ezrin at T567 to restrict ezrin activity to the distal tips of microvilli (Canals *et al*., 2012; Viswanatha *et al*., 2014; Viswanatha *et al*., 2012). Our new observations suggest that EBP50-associated PP1α might not only serve to inactivate ezrin but at the same time to promote the interaction of EBP50 with TMIGD1. TMIGD1 localized at the subapical region of microvilli could thus serve to tether the ezrin - EBP50 - PP1α complex to the membrane in the subapical region of microvilli. In this complex, PP1α would be able to dephosphorylate EBP50-associated ezrin at T567 resulting in ezrin inactivation. The different binding sites in EBP50 for TMIGD1 (PDZ1, AA 14-94 and PDZ2, AA 154 - 234), PP1α (AA 257-259) (Zhang *et al*., 2019) and ezrin (AA 340-358) (Cheng *et al*., 2009) would most likely allow a simultaneous interaction of all three components with EBP50.

This model of a tethering function of TMIGD1 for the ezrin - EBP50 - PP1α complex thereby inactivating ezrin receives further support from two lines of evidence. First, the ezrin - EBP50 interaction is regulated by EBP50 PDZ domain occupation. EBP50 with intact PDZ domains interacts poorly with ezrin and has a rapid turnover at microvilli (Garbett & Bretscher, 2012). Inactivating the two PDZ domains increases the interaction with ezrin and at the same time stabilizes EBP50 at microvilli (Garbett & Bretscher, 2012; Garbett *et al*., 2013). These observations indicated that the binding of EBP50 to a membrane protein through one or both of its PDZ domains destabilizes its interaction with ezrin. Second, the ezrin - EBP50 interaction is regulated by a feedback mechanism. The binding of active ezrin to EBP50 not only stabilizes active EBP50 but also active ezrin itself (Terawaki *et al*., 2006). Dephosphorylation of ezrin at T567 by EBP50-bound PP1α would thus most likely negatively feed back into EBP50’s ability to stabilize active ezrin resulting in a further destabilization of active ezrin. As a result, inactive ezrin would be no longer capable to maintain EBP50 in its open conformation which may result in reduced binding of EBP50 to PDZ domain ligands in the membrane and increased EBP50 turnover. As predicted from these findings, we observed an increase in the turnover of EBP50/S162A in microvilli in the presence of TMIGD1 (Fig. 6). We speculate that the strong TMIGD1 binding to the S162-unphosphorylated PDZ2 of EBP50 could act as allosteric signal that might enhance the negative feedback regulation of ezrin and EBP50. Thus, tethering the ezrin - EBP50 complex by TMIGD1 to the plasma membrane in the subapical region of microvilli, triggered by dephosphorylation of EBP50 at S162 by EBP50-bound PP1α, would contribute to the inactivation of both ezrin and EBP50 and result in increased EBP50 turnover (see Fig. 8 for a model).

A still unexplored question relates to the mechanism by which TMIGD1 is restricted to the lower region of microvilli. The localization of TMIGD1 overlaps with the localization of E3KARP, which localizes to the lower two thirds of microvilli in JEG-3 cells (Garbett *et al*., 2013). Our findings of a direct interaction of TMIGD1 with E3KARP thus opens the possibility that E3KARP is responsible for the localization of TMIGD1 at the subapical region of microvilli. This possibility is appealing since the interaction of TMIGD1 with EBP50 may be very dynamic and regulated by active ezrin as well as by PKC- and PP1α-mediated phosphorylation and dephosphorylation, respectively, of EBP50. A stable localization of TMIGD1 at the subapical region of microvilli combined with a dynamic association with the active ezrin - EBP50 complex could be achieved if TMIGD1 is engaged in trans-homophilic interactions between adjacent microvilli. Our observations indicate that the extracellular domain of TMIGD1 promotes bead aggregation, strongly suggesting trans-homophilic adhesive activity of TMIGD1. Therefore, one could imagine a scenario in which two TMIGD1 molecules present in a trans-interacting dimer interact differently with EBP50 and E3KARP proteins on opposing microvilli. The E3KARP interaction could restrict the dimer to the subapical region whereas the EBP50 interaction could regulate the dynamic interaction of the ezrin - EBP50 complex with the membrane.

Recent observations indicate that microvilli organization is regulated by an intermicrovillar adhesion complex (IMAC) localized at the tips of microvilli (Crawley *et al*., 2014b). This complex consists of the protocadherins cadherin-related family member (CDHR) 2 and CDHR5 which trans-heterophilically interact through their extracellular domains and which are linked to a cytoplasmic protein complex consisting of the PDZ domain protein USH1C/Harmonin, Myosin-7b and ankyrin repeat and SAM domain-containing protein 4B (ANKS4B) (Crawley *et al*., 2014b; Crawley *et al*., 2016; Weck *et al*., 2016). This complex regulates the packing density of microvilli (Crawley *et al*., 2014b). Our observations that TMIGD1 undergoes trans-homophilic interaction, and that TMIGD1 interacts with EBP50 and E3KARP proteins which act as scaffolds for ezrin and PP1α suggest that TMIGD1 forms a second intermicrovillar adhesion complex. We hypothesize that the function of this adhesion complex could be to inactivate ezrin at the subapical region of microvilli thereby restricting ezrin activity to the distal tip region (Fig. 8). Future studies with TMIGD1 knockout mice will be important to analyze the function of TMIGD1 in microvilli formation and dynamics.

## Materials and Methods

### Cell culture and transfections

Caco-2 cells (clone C2BBe1, ATCC-CRL-2102) were grown in DMEM (Sigma-Aldrich (SA) #D5671), 10% FCS, 1% non-essential amino acids (NEAA), 10 µg/ml human transferrin, 2 mM L-glutamine (L-Glu), 100 U/ml penicillin and 100 U/ml streptomycin (Pen/Strep, Biochrom, Berlin). Upon splitting, cells were seeded at 4.5 × 10^3^ cells/cm^2^. Cells were subcultured after reaching approximately 50% of confluency under routine culture conditions to prevent differentiation. T84 cells were grown in DMEM/F-12 1:1 mixture (SA # 6421), 10% FCS, 1% NEAA, 10 µg/ml human transferrin, 2 mM L-Glu, 100 U/ml Pen/Strep. LS174T-W4 cells are human colon-derived epithelial cells stably transfected with a constitutive LKB1 and a tetracycline (tet)-controlled STRAD expression vector in which LKB1 activation can be regulated by tet-induced STRAD expression (Baas *et al*., 2004). Cells were kindly provided by Lucas J. M. Bruurs and Johan L. Bos, Oncode Institute, University Medical Center Utrecht, Utrecht, The Netherlands. Cells were maintained in RPMI 1640 medium (Merck #F1215), 10% tetracycline-free FCS (Clontech/TaKaRa, #631106), 2 mM L-Glu, 100 U/ml Pen/Strep. Expression of STRAD and the concomitant activation of LKB1 was induced by adding doxycycline (1 µg/ml) for at least 16 h. LS174T-W4 cells with a stable KD of EBP50 or E3KARP were generated by lentiviral transduction with either EBP50- or E3KARP shRNAs-expressing plasmids followed by selection in medium containing 3 µg/ml puromycin. EBP50/E3KARP double KD LS174T-W4 cells were generated by lentiviral transduction of stable single KD cells with plasmids encoding shRNAs of the respective homolog followed by selection in medium containing 6 µg/ml puromycin for 3 weeks. Single and double knockdown efficiencies were confirmed by western blot analysis (Suppl. Fig. S2B). JEG-3 cells (ATCC-HTB-36) were grown in MEM (Gibco, #51200-046), 10% FCS, 2 mM L-Glu, 1mM Na-pyruvate (Merck #L0473), 100 U/ml Pen/Strep. For live cell imaging, JEG-3 cells were maintained in phenol red-free MEM (ThermoFisher #51200038) supplemented with the same additives and in addition 20 mM HEPES. JEG-3 cells stably expressing hTMIGD1 from a doxycycline-regulated promoter were generated by transducing JEG-3 cells with a pInducer21-Puro plasmid vector (kindly provided by Dr. T. Weide, Department of Internal Medicine D, Division of Molecular Nephrology, University Hospital Münster, Germany) encoding hTMIGD1. Transduced cells were selected in medium containing 3 µg/ml puromycin (Santa Cruz #sc-108071A) and subcloned by limiting dilution. Inducible expression of TMIGD1 was confirmed by western blot and IF analysis (Suppl. Fig. S3). HEK293T cells (ATCC-CRL-2316) were grown in DMEM containing 10% FCS, 1% NEAA, 2 mM L-Glu, 100 U/ml Pen/Strep.

Transient transfections of cDNAs were performed using Lipofectamine 2000 (ThermoFisher Scientific, #11668-019) and X-Fect (Xfect(tm) Transfection Reagent Clontech/TaKaRa #631318), according to manufacturer’s instructions. Lentiviral particles for the generation of stably transfected cell lines expressing either shRNAs (EBP50, E3KARP) or hTMIGD1 cDNAs were generated by co-transfection of HEK293T cells with the lentiviral vector and the packaging vectors psPAX2 and pMD2.G (kindly provided by Dr Didier Trono, Addgene plasmids 12260 and 12259) in a ratio of 3:2:1 into HEK293T cells. Lentiviral transduction of cells was performed as described (Tuncay *et al*, 2015). The following shRNAs were used: hEBP50/hNHERF1 shRNA in pLKO.1 (5’-CCTAGACTTCAACATCTCCCT -3’, Dharmacon #TRCN0000043736), hE3KARP/hNHERF2 shRNA in pLKO.1 (5’-GATGAACACTTCAAGCGGCTT -3’, Dharmacon #TRCN0000043707).

### Expression vectors

The following constructs were used. Flag-TMIGD1 constructs in pFlag-CMV-1 (N-terminal Flag tag, Sigma-Aldrich, Munich, Germany): hTMIGD1 without signal peptide (AA 30-262) and hTMIGD1 without signal peptide lacking the PDZ domain binding motif (hTMIGD1/Δ5, AA 30-258). TMIGD1 constructs in pInducer21-Puro: hTMIGD1 full length (AA1-262). TMIGD1 constructs in pcDNA3 (Invitrogen): hTMIGD1 full length (AA1-262), hTMIGD1 lacking the PDZ domain binding motif (hTMIGD1/Δ5, AA 1-258), hTMIGD1 lacking the cytoplasmic domain (hTMIGD1/ΔCP, AA 1-241). Flag-tagged TMIGD1-EGFP constructs in pKE1079 (Hartmann *et al*., 2020): EGFP-hTMIGD1 (AA 30-262, EGFP inserted between AA211 and AA212 of TMIGD1); EGFP-TMIGD1-JAM-A (NH_2_-TMIGD1-30-211-EGFP-JAM-A-231-299-COOH). GST-tagged TMIGD1 constructs in pGEX-4T-1 (GE Healthcare): GST-TMIGD1 (hTMIGD1 cytoplasmic tail (AA 242-262); GST-TMIGD1/Δ5: hTMIGD1 cytoplasmic tail lacking the PDZ domain binding motif (AA 242-257). GST-TMIGD1/S258D (hTMIGD1 cytoplasmic tail (AA 242-262_S_258_D); GST-TMIGD1/S260D (hTMIGD1 cytoplasmic tail (AA 242-262_S_260_D). TMIGD1 constructs in yeast-two hybrid vector pBTM116 (Keegan & Cooper, 1996): pBTM116-TMIGD1 (cytoplasmic tail of hTMIGD1, AA 241-262). TMIGD1-Fc fusion constructs in pcDNA3-hIgG): hTMIGD1-Fc (extracellular domain of hTMIGD1, AA 1 - 222).

EBP50/NHERF1 constructs in pKE081myc (N-terminal myc tag, (Ebnet *et al*, 2000): hEBP50 full length (hEBP50, AA 2-358), hEBP50 with mutated PDZ domain 1 (hEBP50_P1M, AA 1-358_G_25_AF_26_A), hEBP50 with mutated PDZ domain 2 (hEBP50_P2M, AA 1-358_G_165_AF_166_A). EBP50 constructs in pEGFP-C3 (Clontech): hEBP50 full length (hEBP50, AA1-358), hEBP50 with mutated PDZ domain 1 (hEBP50_P1M, AA 1-358_G_25_AF_26_A), hEBP50 with mutated PDZ domain 2 (hEBP50_P2M, AA 1-358_G_165_AF_166_A). The same EBP50 constructs containing mutations in the shRNA target site (5’-CCTgGAtTTtAAtATaTCaCT -3’) were generated. hEBP50 with mutations in serine phosphorylation sites: hEBP50_S_162_A (AA 1-358_S_162_A), hEBP50_S_280_AS_302_A (AA 1-358_S_280_AS_302_A), hEBP50_S_339_AS_340_A (AA 1-358_S_339_AS_340_A). GST-hEBP50 in pGEX-6P-2 (GE Healthcare): GST-hEBP50/WT (AA95-168), GST-hEBP50_S_162_A (AA95-168_S_162_A).

E3KARP/NHERF2 constructs in pKE081myc: hE3KARP full length (hE3KARP, AA 2-337), hE3KARP with mutated PDZ domain 1 (hE3KARP_P1M, AA 2-337_G_22_AF_23_A), hE3KARP with mutated PDZ domain 2 (hE3KARP_P2M, AA 2-337_G_162_AF_163_A). E3KARP constructs in pKE1400 (with N-terminal triple-Flag tag): hE3KARP PDZ domains 1 and 2 (Flag-E3KARP/PDZ1-2, AA 2-277). E3KARP constructs in pEGFP-C3 (Clontech): hE3KARP full length (hE3KARP, AA 2-337), hE3KARP with mutated PDZ domain 1 (hE3KARP_P1M, AA 2-337_G_22_AF_23_A), hE3KARP with mutated PDZ domain 2 (hE3KARP_P2M, AA 2-337_G_162_AF_163_A). Identical E3KARP constructs containing mutations in the shRNA target site (5’-GAcGAgCAtTTtAAaCGctTc -3’) were also generated.

Ezrin constructs in pET-28a(+) (N-terminal hepta-His tag, Novagen, Madison, WI): hEzrin wildtype (His-Ezrin/WT, AA 1-586), hEzrin with mutated T567 phosphorylation site (His-Ezrin/T_567_D, AA 1-586_T_567_D). Ezrin constructs in pEGFP-C2 (Clontech): hEzrin wildtype (EGFP-Ezrin/WT, AA 1-586). Ezrin constructs in pcDNA3: hEzrin/Δ3 (hEzrin AA 1-583).

JAM-A and Coxsackie- and Adenodvirus Recptor (CAR) constructs in pcDNA3: mJAM-A (AA 1 - 300), mCAR (AA 1-365). GST-JAM-A, GST-JAM-B and GST-JAM-C constructs containing the cytoplasmic domains of JAM-A, JAM-B and JAM-C fused to GST as well as JAM-B-Fc and JAM-C-Fc fusion constructs containing the extracellular domains of JAM-B and JAM-C have been described before (Ebnet *et al*, 2003).

### Antibodies and reagents

The following antibodies were used in this study: rabbit pAb anti-TMIGD1 (Sigma-Aldrich #HPA021946); rabbit pAb anti-TMIGD1 (proteintech # 27174-1 AP); rabbit pAb anti-EBP50 (ThermoFisher #PAI-090); rabbit pAb anti-NHERF1/EBP50 (Novus #300-536); rabbit pAb anti-SLC9A32 (NHERF2/E3KARP) (Sigma Aldrich # HPA001672); mouse mAb anti-Ezrin (BD-TL #610602), rabbit mAb anti-Phospho-Thr567-Ezrin (CST #3726), rabbit pAb anti-Phospho-Thr567-Ezrin (CST #3141), mouse mAb anti-Villin (SantaCruz #sc-58897); mouse mAb anti-Eps8 (BD-TL #610143); mouse mAb anti-ZO-1 (BD-TL #610966); mouse mAb anti-α-Tubulin (Sigma-Aldrich, clone B-5-1-2, #T5168); mouse mAb anti-Flag M2 (Sigma-Aldrich #F1804); rabbit pAb anti-Flag (Sigma-Aldrich #F7425); goat pAb anti-Myc (SantaCruz #sc-789G); mouse mAb anti-Myc 9E10 (Evan *et al*, 1985). Rabbit anti-TMIGD1 pAbs Affi1662/1663 was generated by immunizing rabbits with a fusion protein consisting of the extracellular domain of hTMIGD1 fused to the Fc region of human IgG, as described previously (Rehder *et al*, 2006). The antibodies were affinity-purified by adsorption at the antigen covalently coupled to cyanogen bromide (CNBr)-activated sepharose beads (Amersham Biosciences Europe, Freiburg, Germany). Antibodies directed against the Fc part were depleted by adsorption at human IgG coupled to CNBr-activated sepharose beads. Affinity-purified antibodies were dialyzed against PBS. Secondary antibodies and fluorophore-conjugated antibodies: Fluorophore-conjugated antibodies for Western blotting: IRDye 800CW Donkey anti-Rabbit IgG (LI-COR Biosciences #926-32213), IRDye 680CW Donkey anti-mouse IgG (LI-COR Biosciences #926-68072). Fluorophore-conjugated secondary antibodies for ICC: Donkey anti-Mouse IgG (H+L) Alexa Fluor 594 (ThermoFisher Scientific #A-21203); Donkey anti-Rabbit IgG (H+L) Alexa Fluor 594 (ThermoFisher Scientific #A-21207); Donkey anti-Rabbit IgG(H+L) Alexa Fluor 488 (ThermoFisher Scientific #A-21206); Donkey anti-Mouse IgG (H+L) Alexa Fluor 488 (Dianova/Jackson ImmunoResearch Europe Ltd #715-545-150); Donkey anti-Mouse IgG (H+L) Alexa Fluor 647 (Dianova/Jackson ImmunoResearch Europe Ltd #715-605-150) -conjugated, highly cross-adsorbed secondary antibodies. The following reagents were used: Doxycycline (SA #D9891), collagen type I (rat tail type 1 collagen, Advanced BioMatrix #5163), lysozyme (SA #L6876), imidazole (Carl Roth #3899), TRITC-Phalloidin (SA #P1951), CytoPainter Phalloidin-iFluor 647 (Abcam #176759), Phorbol 12-myristate 13-acetate (PMA) (SA #P8139), [γ-^32^P]ATP (3000 Ci/mmol, 10 mCi/ml, Hartmann Analytic GmbH, Braunschweig, Germany, # SCP-301), PKCα (Eurofins Discovery, Cell L’Evescault, France, #14-484), PKC lipid activator (MerckMillipore #20-133), PP1α (Eurofins #14-595).

### Yeast Two-hybrid Screen

Yeast two-hybrid screening experiments were performed essentially as described (Ebnet *et al*., 2000). Briefly, the Saccharomyces cerevisiae reporter strain L40 expressing a fusion protein between LexA and the cytoplasmic tail of TMIGD1 (AA 241–262) was transformed with 250 µg of DNA derived from a day 9.5/10.5 mouse embryo cDNA library (Hollenberg *et al*, 1995) according to the method of Schiestl and Gietz (Schiestl & Gietz, 1989). The transformants were grown for 16 h in liquid selective medium lacking tryptophan, leucine (SD-TL) to maintain selection for the bait and the library plasmid, then plated onto synthetic medium lacking tryptophan, histidine, uracil, leucine, and lysine (SD-THULL) in the presence of 1 mM 3-aminotriazole. After 3 days at 30 °C, large colonies were picked and grown for additional three days on the same selective medium. Plasmid DNA was isolated from growing colonies using a commercial yeast plasmid isolation kit (DualsystemsBiotech, Schlieren, Switzerland). To segregate the bait plasmid from the library plasmid, yeast DNA was transformed into E. coli HB101, and the transformants were grown on M9 minimal medium lacking leucine. Plasmid DNA was then isolated from E. coli HB101 followed by sequencing to determine the nucleotide sequence of the inserts.

### Immunoprecipitation and Western blot analysis

For immunoprecipitations, cells were lysed in lysis buffer (25 mM TrisHCl, pH 7.4, 1% (v/v) Nonidet P-40 (NP-40, AppliChem, Darmstadt, Germany), 150 mM NaCl, protease inhibitors (Complete Protease Inhibitor Cocktail; Roche, Indianapolis, IN), 5% glycerol ()AppliChem #A2926) and phosphatase inhibitors (PhosSTOP™, Roche, Indianapolis, IN), 2 mM sodium orthovanadate) for 20 min with overhead rotation at 4°C followed by centrifugation (15.000 rpm, 20 min at 4°C). Postnuclear supernatants were incubated with 3 μg of antibodies coupled to protein A– or protein G–Sepharose beads (GE Healthcare, Solingen, Germany) for 4 h at 4°C. Beads were washed five times with lysis buffer, bound proteins were eluted by boiling in 3x SDS-sample buffer/150 mM DTT. Eluted proteins were separated by SDS–PAGE and analyzed by Western blotting with near-infrared fluorescence detection (Odyssey Infrared Imaging System Application Software Version 3.0 and IRDye 800CW-conjugated antibodies; LI-COR Biosciences, Bad Homburg, Germany).

### Purification of recombinant His-tagged proteins

(His)_6_-tagged ezrin and (His)_6_-tagged ezrin T567D was expressed in E. coli cells (strain BL21(DE3)pLysS). Transformed bacteria were grown to an OD_600_ of 0.6 at 37°C, expression of recombinant proteins was induced by incubating cells with isopropyl β-D-1-thiogalactopyranoside (IPTG, 1 mM) for 4 to 5 h at 37 °C. Cells were harvested by centrifugation (4000 ×g, 15 min) and resuspended in ice-cold lysis buffer (50 mM NaH_2_PO_4_, pH 8, 300 mM NaCl, 10 mM imidazole, 10 mM EDTA, 10 mM β-mercaptoethanol, 1 mM PMSF). Cells were lysed by 3 freeze-thaw (liquid nitrogen – 37°C waterbath) cycles, incubation with 1 mg/ml lysozyme for 30 min on ice followed by sonication (directional cycle 50, output control 5 for 35s, 3x). After centrifugation (50.000 ×g, 45 min), the supernatant was applied to a Ni–NTA-agarose column (SA #P6611) pre-equilibrated with lysis buffer and incubated for 1 h at 4°C under gentle rotation. After washing with washing buffer (50 mM NaH_2_PO_4_, pH 8, 300 mM NaCl, 20 mM imidazole) His-tagged ezrin constructs were eluted with elution buffer (50 mM NaH_2_PO_4_, 300 mM NaCl, 250 mM imidazole) and dialyzed against PBS. Protein solutions were adjusted to 50% (v/v) glycerol and stored at -20 °C. Purified proteins were analyzed by SDS-PAGE and Coomassie Brilliant Blue staining.

### GST pulldown experiments

*In vitro* binding experiments were performed with recombinant GST fusion proteins purified from *E*.*coli* and immobilized on glutathione-Sepharose 4B beads (Life Technologies #17-0756-01). Purification of GST fusion proteins was performed as described (Ebnet *et al*., 2000). For protein interaction experiments the putative partner protein (prey) was expressed in HEK293T cells by transient transfection. Cells were lysed as described for immunoprecipitations. Lysates were incubated with 3 µg of immobilized GST fusion protein for 2 h at 4°C under constant agitation. After 5 washing steps in lysis buffer, bound proteins were eluted by boiling for 5 min in SDS sample buffer, subjected to SDS-PAGE and analyzed by western blotting using prey-specific antibodies.

### *In vitro* phosphorylation and dephosphorylation assays

Phosphorylation studies were performed essentially as described before (Iden *et al*, 2012). Briefly, GST fusion proteins were coupled to glutathione sepharose (GE Healthcare, Freiburg, Germany) using 1x buffer B (10 mM HEPES-NaOH pH 7.4, 100 mM KCl, 1 mM MgCl_2_, 0.1% Triton X-100) and subjected to *in vitro*-kinase reactions with recombinant PKCα. Assays were carried out with 10 ng recombinant enzyme and 5 µCi γ-[^32^P]-labeled ATP in PKCα kinase buffer (20 mM HEPES pH 7.4, 700 µM CaCl_2_, 25 mM MgCl_2_, 0.25 mM cold ATP) for 30 min at 30°C in the presence of 1:10 diluted PKC lipid activator (10x lipid activator: 0.5 mg/ml phosphatidylserine, 50 µg/ml 1-stearoyl-2-linoleoyl-sn-glycerol, 50 µg/ml 1-oleoyl-2-acetyl-sn-glycerol, 0,15 % Triton X-100, 1 mM DTT, 2 mM CaCl_2_, 20 mM MOPS, pH 7.2). For *in vitro* dephosphorylation assays, the phosphorylated GST fusion proteins were washed in buffer B, then incubated for 30 min at RT with 2 U/sample recombinant PP1α (Eurofins) in PP1α reaction buffer (57 mM HEPES pH 7.2, 10 mM MnCl_2_, 0.167 mM DTT, 0.83% (v/v) glycerol, 0.0167 % BSA, 0.002% Brij 35). The GST fusion proteins were washed and subsequently eluted from the beads by boiling in SDS sample buffer. Eluted proteins were separated by SDS-PAGE and analyzed by autoradiography.

### Cell aggregation assays

Cell aggregation assays were performed essentially as described previously (Brinkmann *et al*, 2016). Briefly, HEK293T cells were transiently transfected with expression vectors encoding JAM-A, CAR or TMIGD1. After 24 h cells were harvested, resuspend in HBSS and then adjusted to 5 × 10^5^ cells/ml. 3 ml of cell suspension were added to a 30-mm BSA-coated 6-well tissue culture dishes and incubated on a horizontal shaker at 40 rpm for 45 min at 37°C. Cells were fixed by addition of 500 µl 25% glutaraldehyde and incubation on ice for 30 min. Fixed cells were analyzed by microscopic analysis at 10x magnification. Five to ten pictures were analyzed for each condition. The number of particles was counted using ImageJ software, particles larger than 256 µm^2^ were considered as aggregates. Data are presented as means ± standard error (SEM) from five independent experiments. P-values: *P<0.05, ***P<0.001 and ****P<0.0001.

### Bead aggregation assays

Bead aggregation assays were performed essentially as described (Emond *et al*, 2011). Briefly, equal amounts of Fc-fusion proteins consisting of the extracellular domains of TMIGD1 (TMIGD1-Fc), JAM-B (JAM-B-Fc) or JAM-C (JAM-C-Fc) fused C-terminally to the Fc-part of human IgG were coupled to red fluorescent, protein A-coated beads (BNF-Starch-redF/ProteinA, 100 nm, micromod Partikeltechnologie GmbH, Rostock, Germany, #64-20-102). Beads were washed three times in aggregation buffer (50 mM Tris-HCl (pH 7.4), 100 mM NaCl, 10 mM KCl, 0.2% BSA) to remove unbound Fc-fusion proteins. After trituration by vigorous pipetting and vortexing, beads were resuspended in aggregation buffer transferred to 15-ml conical tubes and incubated for 30 min at RT °C under constant shaking (180 rpm) Beads were carefully transferred to microscopic slides. Fluorescent images were collected on a microscope (LSM 800) with a 10x Plan-Apochromat x 63/1.4 oil objective. Aggregate formation was analyzed by measuring the total area of fluorescence within a given field of view using ImageJ software. At least 25 fields of view were analyzed per experiment (three independent experiments). Statistical analysis was performed using Ordinary one-way ANOVA with Dunnett’s multiple comparison, data are presented as means ± standard error (SEM). P-value: ****P<0.0001.

### Immunofluorescence microscopy

For immunofluorescence microscopy, cells were grown on collagen-coated glass slides or collagen-coated transwell polycarbonate membrane filters (0.4 μm pore size, Corning, Amsterdam, The Netherlands). Cells were washed with PBS and fixed with 4% paraformaldehyde (PFA, SA) for 7 min. To detect intracellular proteins, PFA-fixed cells were incubated with PBS containing 0.2% Triton X-100 for 15 min. Cells were washed with 100 mM glycine in PBS, blocked for 1h in blocking buffer (PBS, 10% FCS, 0.2% Triton X-100, 0.05% Tween-20, 0.02% BSA) and then incubated with primary antibodies in blocking buffer for 1 h at room temperature (RT) or overnight at 4 °C. After incubation, cells were washed three times with PBS and incubated with fluorochrome (AlexaFluor488, AlexaFluor594 and AlexaFluor647)-conjugated, highly cross-adsorbed secondary antibodies (Invitrogen) for 2 hrs at RT protected from light. F-Actin was stained using phalloidin-conjugates (TRITC and cytoPainter iFluor-647, DNA was stained with 4,6-diamidino-2-phenylindole (DAPI, Sigma-Aldrich). Samples were washed three times with PBS and mounted in fluorescence mounting medium (Mowiol 4-88, Sigma Aldrich). Immunofluorescence microscopy was performed using the confocal microscopes LSM 780 and LSM 800 Airyscan (both from Carl Zeiss, Jena, Germany) equipped with the objectives Plan-Apochromat x 63/1.4 oil differential interference contrast (Carl Zeiss). Image processing and quantification was performed using ImageJ, Zen 2 (Blue Edition, Carl Zeiss) and Imaris (Bitplane, Version 9.1.2) software.

For quantification of TMIGD1 recruitment by EBP50 and/or E3KARP to polarized caps of LS174T-W4 cells, a total number of at least 92 and a maximum of 142 cells per condition (3 independent experiments) was analyzed. Only cells containing a polarized brush border (polarized ezrin or Eps8 signals after doxycycline-treatment) and successfully transfected with TMIGD1 were included in the analysis. Statistical analysis was performed using unpaired student’s t test, data is plotted as means ± standard deviation (SD). P-values: *P<0.05, ****P<0.001.

### FRAP experiments

For FRAP experiments, JEG-3 cells grown on collagen-coated glass bottom slides (Ibidi #80827) were analyzed by time-lapse microscopy on a confocal-laser scanning microscope (Carl Zeiss, Jena Germany, LSM 800 Airyscan) in a 37°C chamber. FRAP measurements and quantifications were performed essentially as described before (Kondadi *et al*, 2020). A square region of 3.7 × 3.7 µm was bleached using 100% laser power of URGB Diodes at 488 nm (Diode laser 488 nm, 10 mW). To monitor fluorescence recovery over time 5 pre-bleach images and 145 - 195 post-bleach images of a given region of interest (ROI) were acquired. For FRAP quantification three ROIs were defined: ROI_1_ (region with no microvilli, used to perform background subtraction), ROI_2_ (region with microvilli where photobleaching was performed), ROI_3_ (region with microvilli where photobleaching was not performed, used to obtain the correction factor for the amount of bleaching during monitoring). Photobleach correction was performed by dividing the background-subtracted intensity values obtained at ROI_2_ (ROI_2_ - ROI_1_) by those obtained at ROI_3_ (ROI_3_ - ROI_1_). Normalization of ROIs was performed with ImageJ software (Fiji) using the FRAP Norm plugin (https://imagejdocu.tudor.lu/doku.php?id=plugin:analysis:frap_normalization:start.) which uses the normalization method outlined in (Phair *et al*, 2004). Normalized FRAP recovery values were fitted using GraphPad Prism 6 by nonlinear regression two phase association model. FRAP graphs represent fitted curves with means ± standard deviation (SD). First and second association t_1/2_ recovery values were obtained when the curves were fitted by nonlinear regression two-phase association model. For each condition a total number of 50 to 143 cells obtained from 3 independent experiments were analyzed. Statistical analysis of differences between recovery curves of cells without and cells with TMIGD1 expression was performed using Two-way Repeated Measurements ANOVA. *P<0.05, ***P<0.001.

### Intestinal organoids

Organoids derived from intestinal crypts were generated essentially as described (Mosa *et al*, 2018). Briefly, after mechanical removal of villi, intestinal tissue was cut into small tissue pieces (2 – 4 cm) and incubated in PBS/2mM EDTA for 30 min on ice. The material was passed through a 70 µm strainer, followed by centrifugation (140xg, 5 min, 4°C). The pellet was resupended in AD-DF+++ medium (Advanced DMEM/F12 (Gibco, #12634-010), 2 mM L-Glu (Lonza, #17-602E, 100 U/ml Pen/Strep (Lonza, #17-602E), 10 mM HEPES (Sigma, #H0887) and resupended in ice-cold matrigel. 50-µl aliquots of the matrigel suspension were transferred to pre-warmed 24-well plates, allowed to solidify, then covered with complete growth medium (80 % AD-DF+++, 10 % RspondI conditioned medium, 10 % Noggin conditioned medium, 0.5 µg/ml mEGF (Gibco, #PMG8043), 0.1 mg/ml Primocin (InvivoGen, #ant-pm-0.5), 1X B-27 supplement (Gibco, #175044), 1.25 mM N-Acetyl-L-Cysteine (Sigma, #A9165-5G)). Organoids were cultured for 7 days with medium changes every 2 to 3 days. For continuous culture, organoids were mechanically disrupted by vigorous pipetting to generate single crypts, then resuspended in matrigel and transferred to new tissue culture dishes as described above. Organoids were maintained as a polyclonal pool.

For immunofluorescence staining crypts were grown for 3 days in single matrigel droplets on 15 µ-slide 8 well glass bottom ibidi slides (#80827, Ibidi). Prior to fixation the matrigel was dissolved by incubation on ice in cell-recovery solution (#354253, Corning) for approx. 1 h. Organoids were fixed in 4% PFA/PBS for 1h at room temperature, washed with PBS and incubated in quenching solution (130 mM NaCl, 100 mM glycine, 7 mM Na_2_HPO_4_ and 3.5 mM NaH_2_PO_4_; 30 min). After permeabilization (0.5% Triton X-100 in PBS for 1h) and blocking (PBS, 0.02% BSA, 0.05% Tween20, 0.05% Triton X-100, 5% goat serum, 5% donkey serum, at least 2 h) organoids were incubated with primary antibodies in blocking buffer at 4°C overnight followed by incubation with fluorophore-conjugated secondary antibodies for 2 h at RT. Organoids were imaged with a Zeiss LSM800 using the Airyscan mode and a 63 x oil immersion objective (numeric aperture, 1.4).

### Electron microscopy

Electron microscopy was performed with Caco-2_BBe_ cells grown on polycarbonate filters for 22 d to induce differentiation. For pre-embedding immunogold labelling, cells were fixed with 2% paraformaldehyde (Polysciences, Warrington PA, USA) in PBS (pH7.4). Cells were incubated with rabbit anti-TMIGD1 pAb, followed by incubation with 15-nm gold particle-conjugated protein A (Department of Cell Biology, University Medical Center, Utrecht, The Netherlands). After fixation with 2% glutaraledhyde (Polysciences) in PBS (pH7.4), samples were counterstained with osmiumtetroxide (Polysciences) and embedded in epoxy resin (Agar Scientific, Essex, UK). 60-nm ultrathin sections were generated using an ultramicrotome (Leica-UC6 ultramicrotome, Vienna, Austria). For immunogold labelling of ultrathin cryosections, cells were fixed with 2% paraformaldehyde, 0.2% glutaraldehyde in PBS (pH 7.4) and processed for immuno-labelling electron microscopy according to the method of Tokuyasu (Tokuyasu, 1980). 60-nm ultrathin sections (Leica-UC6 ultramicrotome) were labelled with rabbit anti-TMIGD1 pAb as described above. All samples were analyzed at 80 kV on a FEI-Tecnai 12 electron microscope (FEI, Eindhoven, The Netherlands). Selected areas were documented with Veleta 4k CCD camera (EMSIS GmbH, Münster, Germany). Analysis of TMIGD1 immunogold-labelling was performed on three independent sections and a total of 340 individual cells. Of these cells, 23 (6.8%) were positive for gold particles.

## Acknowledgements

We gratefully acknowledge the help of Drs Lucas J. M. Bruurs, Johan L. Bos and Hans Clevers (Oncode Institute, University Medical Center Utrecht, Utrecht, The Netherlands) for providing us LS174T:W4 cells. We thank Dr. H.-J. Kreienkamp (Institute for Human Genetics, University Medical Center Hamburg-Eppendorf, Hamburg, Germany) for providing us with the NHERF1-GFP plasmid vector. We also thank Dr. Thomas Weide (Department of Internal Medicine D, Division of Molecular Nephrology, University Hospital Münster, Germany) for providing us with the pInducer21-Puro plasmid vector.

## Author contributions

C.H. and K.E. designed and conceived the study. C.H., E.-M.T., B.E.M., D.P., S.L., L.G., F.B. and M.G.-E. performed experiments. C.H., L.G., E.W., T.Z., M.A.S., V.G. and K.E. analyzed the data. C.H. and K.E. wrote the manuscript.

### Abbreviations

AA: amino acid
E3KARP: NHE3 kinase A regulatory protein
EBP50: ERM-binding phosphoprotein of 50 kDa
FRAP: fluorescence recovery after photobleaching
IF: immunofluorescence
JAM-A: Junctional Adhesion Molecule-A
NHERF: Na^+^/H^+^ exchange regulatory cofactor
PBM: PDZ domain-binding motif
PDZ: Postsynaptic density 95 - Discs Large - Zonula occludens-1
TMIGD1: Transmembrane and immunoglobulin domain-containing protein 1

## Funding resources

This work was supported by grants from the Deutsche Forschungsgemeinschaft (EB 160/5-1; EB 160/8-1; EXC 1003-CiMIC) and from the Medical Faculty of the University of Münster (IZKF Eb2/020/14).

## Expanded View / Supplemental Figure Legends

**Suppl. Figure S1:**
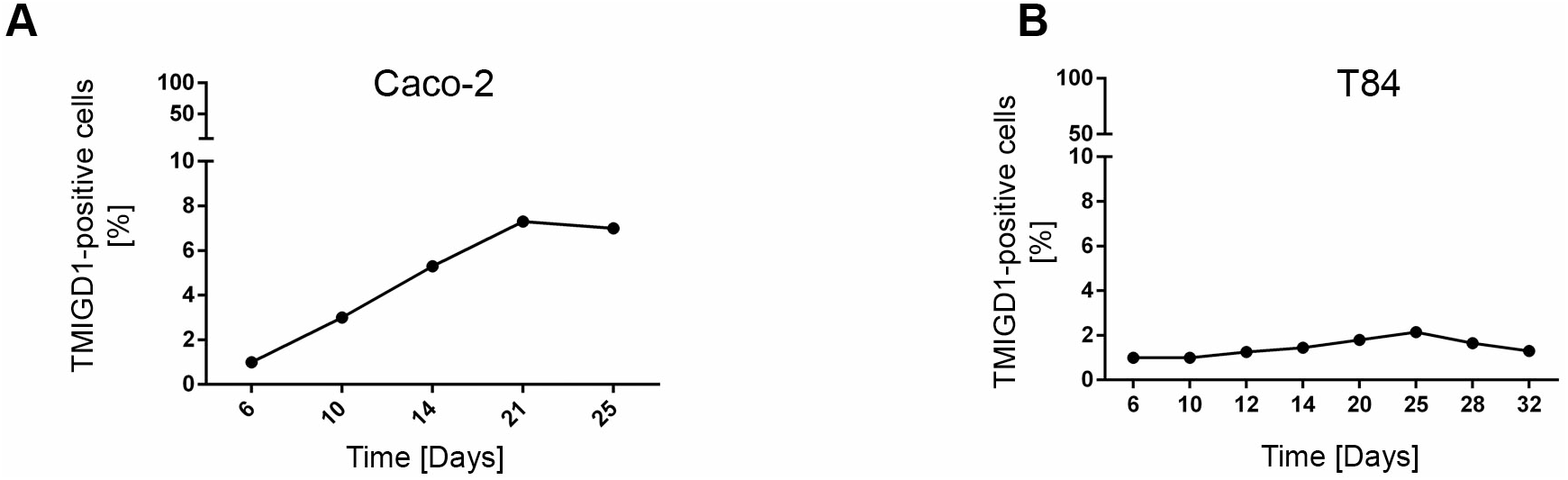
Statistical analysis of TMIGD1 expression in intestinal epithelial cell lines. Caco-2_Bbe1_ cells (left panel) and T84 cells (right panel) were cultured on 0.4 µm polycarbonate filters to allow differentiation. At the indicated time points cells were stained with TMIGD1 antibodies. TMIGD1-positive cells were counted by visual inspection. TMIGD1-positive Caco-2_Bbe1_ cells were quantified in three independent experiments in which at least three fields of view each containing between 81 and 122 cells were analyzed (average fraction of TMIGD1-positive cells at day 24 post confluency: 6.2%, 5.7%, 5.9%).

**Suppl. Figure S2:**
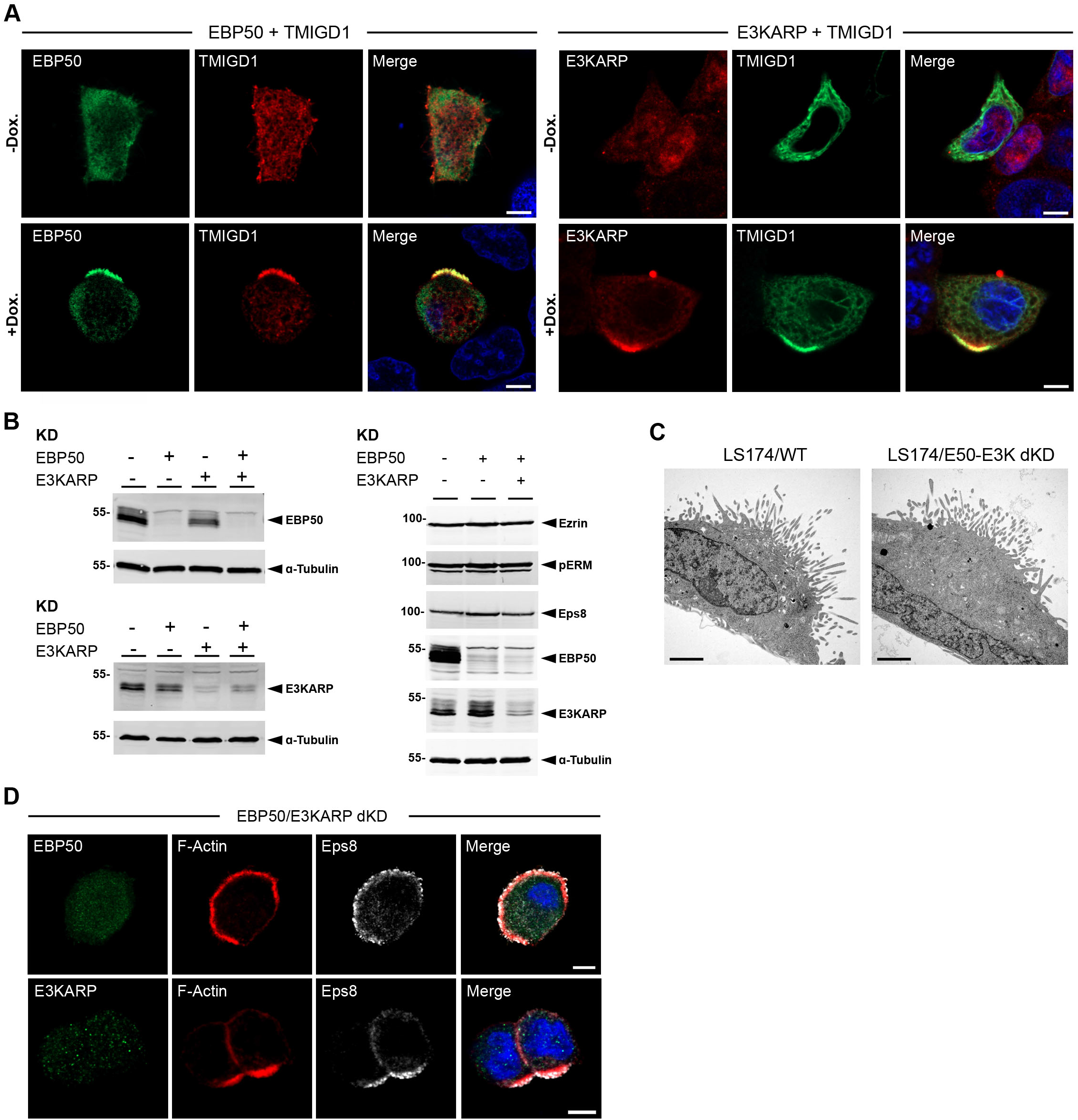
Characterization of LS174T-W4 cells. (**A**) EBP50 and E3KARP co-localize with TMIGD1 at polarized brush borders of LS174T-W4 cells. Cells transfected with untagged TMIGD1 and either EGFP-EBP50 or Myc-E3KARP as indicated were either left uninduced (-Dox.) or were induced (+Dox.) to develop a brush border. Cells were stained with antibodies against TMIGD1 and the Myc-tag and analyzed by IF microscopy. EGFP-EBP50 was detected by its GFP fluorescence. Scale bars: 5 µm. (**B**) EBP50 KD LS174T-W4 cells, E3KARP KD LS174T-W4 cells and EBP50/E3KARP double KD LS174T-W4 cells were analyzed by Western blotting for the expression of EBP50 and E3KARP (left panels) and for the expression of microvilli components ezrin, P-T567-ezrin (pERM) and Eps8 (right panels). (**C**) LS174T-W4 cells (LS174/WT) and EBP50/E3KARP double KD LS174T-W4 cells (LS174/E50-E3K dKD) were analyzed by EM. Note that EBP50/E3KARP double KD LS174T-W4 cells form a microvilli-containing brush border. Scale bars: 2 µm. (**D**) EBP50/E3KARP double KD LS174T-W4 cells induced with doxycycline were stained for F-actin and Eps8 as indicated. Note that F-actin and Eps8 are enriched at polarized brush borders. Scale bars: 5 µm.

**Suppl. Figure S3:**
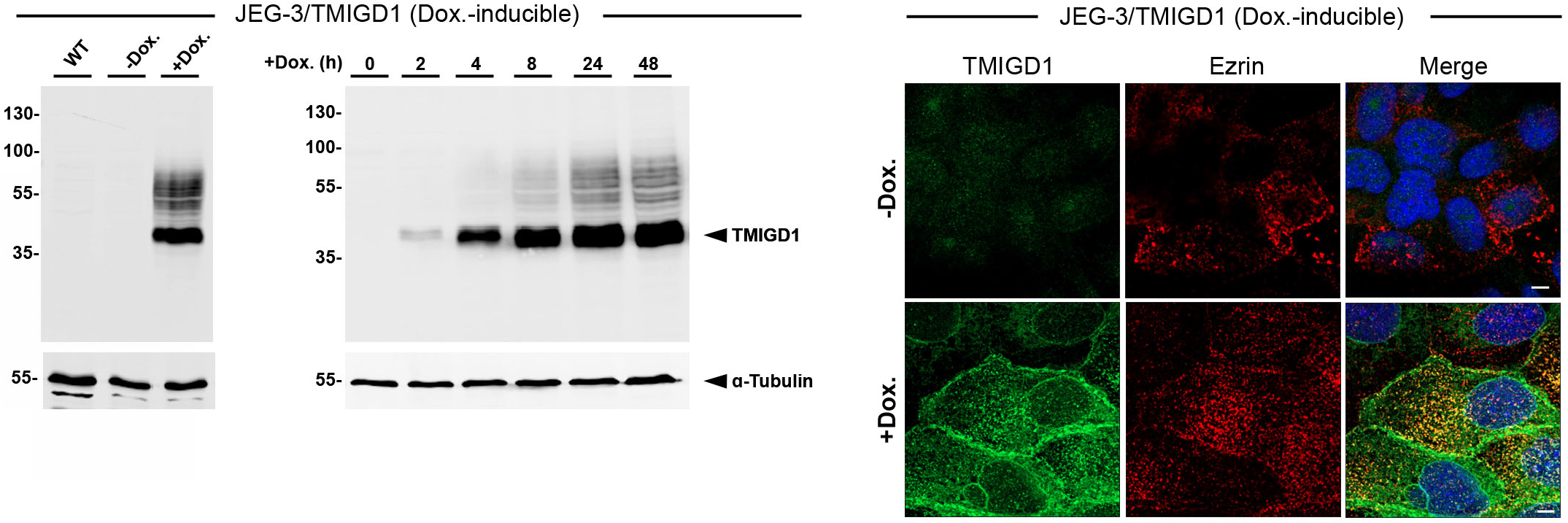
Characterization of JEG-3 cells with inducible expression of TMIGD1. Left panel: Western blot analysis of ectopic TMIGD1 expression in JEG-3 cells. Left blot: JEG-3 cells stably transfected with a lentiviral expression vector allowing inducible expression of TMIGD1 (pInducer21) were treated with doxycycline (Dox.) for 24 h and analyzed with antibodies against TMIGD1. Right Blot: Cells were treated for different time periods with doxycycline as indicated and analyzed with antibodies against TMIGD1. Right panel: IF analysis of TMIGD1 localization in JEG-3 cells. Cells were either left uninduced (-Dox.) or were induced (+Dox.) to express TMIGD1. Cells were stained with antibodies against TMIGD1 and ezrin as indicated. Scale bars: 5 µm.

